# Evolutionary Adaptations of TRPA1 Thermosensitivity and Skin Thermoregulation in Vertebrates

**DOI:** 10.1101/2025.03.31.646411

**Authors:** Gabriel E. Bertolesi, Neda Heshami, Sarah McFarlane

**Author notes:** Correspondence should be addressed to Gabriel E. Bertolesi. NH:, SM.

## Abstract

Altering skin color and reflectance is crucial for temperature regulation in poikilothermic vertebrates, while less so in homeotherms like birds and mammals, which evolved feathers, fur, and other insulation for endothermy. Heat-sensing in vertebrates relies primarily on Transient Receptor Potential (TRP) channels, with certain channels (TRPA1) shifting thermosensitivity over evolution and others retaining heat sensitivity (TRPV1). Exploration of a role for TRP channels in skin physiology has largely focused on human pigmentation and overlooked the evolution of different thermoregulatory structures in the integument of distinct vertebrates. For instance, colour/reflector pigment cells in ectotherms, fur and feathers in endotherms, hairless skin in hominids, and blubber in marine mammals. Therefore, we investigated whether a TRP channel mediates skin darkening induced by heat in the ectotherm *Xenopus laevis* and then explored the evolution of TRPA1 thermal sensitivity and its link with skin physiology. We find Trpa1 mediates heat-induced melanosome dispersion, darkening skin under warmer conditions. In contrast, TRPA1 is known to mediate cold sensation in rodents and UV-induced tanning in humans, leading us to investigate the co-evolution of TRPA1 and skin thermoregulation. Our findings reveal TRPA1 is a heat sensor in ectotherms with uncovered integuments. In mammals, we suggest TRPA1 was thermally insensitive in Euarchontoglires but became cold-sensitive in several rodent lineages. TRPA1 shows reduced selection pressure for thermosensitivity in aquatic mammals (manatees, whales) that depend on blubber for insulation as compared to their terrestrial relatives. These findings emphasize adaptive evolution of TRPA1 in vertebrates, linking thermal sensitivity to the evolution of skin physiology.

## INTRODUCTION

Skin color and/or reflectance alterations play a vital role in regulating body temperature to sustain life (Cuthill et al., 2017; Stuart-Fox et al., 2017). These mechanisms are crucial for survival of poikilotherm vertebrates, but are less relevant for birds and mammals, where feathers and fur in their integument emerged as integral traits for insulation during the evolution of endothermy (Lovegrove, 2017). Members of the Transient Receptor Potential (TRP) non-selective cation channel family are considered the classical thermosensors of vertebrates, although their range of temperature sensitivity differs drastically between lineages (Hoffstaetter et al., 2018; Zhang et al., 2022). This is particularly true of TRPA1 that is a heat sensor in poikilotherm vertebrates [ea. amphibians and reptiles (Cordero-Morales et al., 2011; Saito et al., 2012, 2016)], a cold sensor in rodents, and temperature insensitive in primates (Chen et al., 2013; Kwan et al., 2006; Zappia et al., 2017). Unknown is the role of TRPA1 in regulating skin colour and/or reflectance alterations, and the extent to which the physiology of skin pigmentation and the evolutionary divergence of TRPA1 temperature sensitivity are linked.

Variations in color change and reflectance are generated by the movement of pigmented or reflective organelles located in dermal chromatophores. Eight types of chromatophores containing absorptive or reflective pigments are present in vertebrates (Schartl et al., 2016). *Xenopus laevis* tadpoles offer an advantageous model for studying skin colour change, as the melanin pigment containing melanophores are the main mediators of color change, through organelle dispersion to cause skin darkening and aggregation to induce skin lightening (Bagnara, 1960; Bertolesi et al., 2022). In this species, Trpm8 in melanophores senses cold to induce pigment aggregation and skin lightening (Malik et al., 2023). TRPM8, as a cold sensor, is relatively conserved in vertebrates, though it underwent significant evolutionary change in thermal activity during the transition of organisms from water to land (Lu et al., 2022). Unknown is whether melanophores express a heat sensor to induce melanosome dispersion and skin darkening under warmer conditions in order to counteract the skin lightening induced by cold through Trpm8.

Trpv1 and Trpa1 are both candidates for the heat sensor to regulate skin colour in poikilothermic vertebrates. These channels are often co-expressed, as per in dorsal root ganglion neurons (Masuoka et al., 2017). Depending on the species, however, the temperature sensitivity of TRPA1 differs. For instance, TRPA1 is a cold sensor in mouse, rat and other mammals (Kwan et al., 2006; Story et al., 2003; Zhang et al., 2022), and a heat sensor in amphibians and reptiles (Saito et al., 2012, 2016). Human TRPA1 thermal sensitivity is controversial (reviewed by (Laursen et al., 2015; Tominaga & Iwata, 2025; Zhang et al., 2022)). However, when identical experimental conditions are used to determine sensitivity, rodent TRPA1 functions as a cold sensor, reptiles as a heat sensor and human TRPA1 is insensitive (Chen et al., 2013; Cordero-Morales et al., 2011). Trpa1 actually forms a dimer with Trpv1 to mediate thermal sensitivity in a group of embryonic turtle terminal sensory neurons (Patil et al., 2020) to control heat-induced thermotaxic behavior (Ye et al., 2021). These two channels are co-expressed in melanocytes of human skin (Jia et al., 2021), the mammalian equivalent of poikilothermic melanophores. Ultraviolet (UV) light, however, rather than temperature activates TRPA1 to darken the skin through melanin synthesis (Bellono & Oancea, 2013; Wu et al., 2023). This finding suggests that evolutionary changes in TRPA1 altered its physiological role in skin pigmentation. The conservation of TRPV1 as a heat sensor throughout vertebrate evolution (Hoffstaetter et al., 2018) may also have influenced the function of TRPA1 as a thermosensor in pigmentary physiology. We found that warm temperatures induce skin darkening in *Xenopus laevis* tadpoles and examined the expression of Trpv family members and Trpa1 in the skin and melanophores to identify the channel responsible for heat sensing.

While thermoregulation is a critical physiological function shared between melanin-mediated coloration and TRPA1, significant evolutionary events likely altered their respective roles in maintaining thermal homeostasis. First, by conquering the land, terrestrial ectothermic reptiles gained the ability to adjust their skin pigmentation to regulate body temperature through the reflection or absorption of irradiated energy (Moreno Azócar et al., 2020; Stuart-Fox et al., 2017). Second, in extinct marine reptiles, the melanisation data from fossilized skin suggest a contribution of pigmentation to the adaptation to cold-water habitats (Lindgren et al., 2014, 2015; McNamara et al., 2021). Pigmentation mechanisms became less crucial in homeotherms with feathers or fur, as the advent of endothermy sculpted their skin physiology. Hominids, however, lost body hair and gained sweat glands during the evolution of bipedalism as a more efficient system to dissipate heat (Dávid-Barrett & Dunbar, 2016; Ruxton & Wilkinson, 2011). Interestingly, in humans, melanin synthesis in skin melanocytes is strongly induced by UV light rather than temperature, a pigmentation physiological response that requires TRPA1 activation (Bellono et al., 2013; Maglie et al., 2021; Wu et al., 2023). Whether changes in the thermal activation of TRPA1 in vertebrate melanophores/melanocytes are linked to evolutionary adaptations in thermoregulation or skin physiology remains unknown.

To explore a possible link, we analyzed for the first time a thermosensor role for Trpa1 in skin pigmentation of ectotherms, turning to *Xenopus* laevis as we and others have shown it is an excellent model to identify the molecular mechanisms that underlie the regulation of skin pigmentation (Bertolesi et al., 2022; El Mir et al., 2025). Second, we conducted an evolutionary study, comparing the thermal sensitivity of TRPA1 in related extant species of chordates as identified by electrophysiological data from the literature, with the TRPA1 mutation rate in known temperature sensing regions of the protein. Evolutionary pressures shaped skin structures to optimize thermoregulation, selecting fur and feathers for thermal insulation, energy radiation absorption/reflection in skin pigment cells in organisms with uncovered integument, thick layers of blubber in marine mammals, and sweat glands in hairless human skin. We were particularly interested in the TRPA1 mutation rate in marine mammals, where thermoregulation primarily depends on cardiovascular adjustments rather than skin pigmentation (Favilla et al., 2022). Mammals re-entered the marine environment at least seven times throughout evolution, with extant species originating from five of these events (Uhen, 2007). Notably, the anatomical and physiological changes in the integument differ for each instance of re-entry, reflecting evolutionary adaptations. In general, early adapted species [ea. Sirenia (manatees and dugongs) and Cetaceans (whales and dolphins)] have hairless and thick skin with a blubber layer, the latter essential for thermoregulation. Blubber acts as both a thermal barrier and being highly vascularized allows adjustment of blood flow to the skin and extremities to conserve or dissipate heat as needed (Favilla et al., 2022; Khudyakov et al., 2022). A more recently adapted lineage, the pinnipeds (seals and sea lion), relies on both blubber and fur and behaviour for thermoregulation, while polar bears possess a dense haired fur for thermoregulation (Carolan et al., 2025; Espregueira Themudo et al., 2020; Khamas et al., 2012; Le Duc et al., 2022; Vetter et al., 2001). These marine mammals provide an excellent test to ask whether the evolutionary changes in the thermoregulatory mechanisms between marine mammals and their terrestrial relatives, whose thermoregulatory mechanisms did not undergo large evolutionary change, are associated with differences in selection pressure on the thermosensor region of their respective TRPA1 proteins.

We show that heat disperses melanosomes, darkening the skin of *Xenopus laevis* and melanophores *in vitro* in a manner that we find through pharmacology and siRNA knockdown depends on Trpa1 as the heat sensor. Our investigation of the evolution of TRPA1 thermosensation in chordates, including *Xenopus*, indicates that TRPA1 was initially a heat sensor. Cold thermal sensitivity is present only in certain rodent lineages [Myomorphs (Mice and rats) and Castorimorphs (Bivers and gophers)], which possess an essential amino acid (G878) absent in all remaining species of Euarchontoglires [ea. primates, Lagomorphs (rabbits, hares, and pikas), and Sciuromorphs (squirrels)], in which TRPA1 is thermally insensitive. Interestingly, our comparative analysis of TRPA1 in mammals that returned to a marine habitat during evolution shows that only the Sirenians and Cetaceans, which adapted by losing hair and developing thick blubber, exhibit a higher mutational rate in the protein region responsible for thermoregulation as compared to their terrestrial relatives, suggestive of less thermal positive selection pressure. Collectively, our data reveal a functional role for TRPA1 both in skin pigmentation and more broadly in the various thermoregulatory adaptations that appeared throughout chordate evolution, from the development of fur and feathers to adaptations for marine environments.

## RESULTS

### Determination of an ecologically relevant heat condition for *Xenopus laevis* tadpoles

To determine an ecologically relevant heating paradigm for *Xenopus laevis*, we analyzed the average temperatures recorded over the last 30 years at three national parks in southern Africa (Etosha in Namibia, Kruger in South Africa, and Hwange in Zimbabwe). These national parks are located in regions where *Xenopus laevis* is native (Furman et al., 2015). We examined the temperatures during November and December, which exhibit a large temperature fluctuation between cold (mean daily minimum and cold nights) and hot (mean daily maximum and hot days) conditions (Table 1). The daily temperature fluctuations during these two months oscillate between 37 °C (hot) and 16 °C (cold) (Table 1). However, since *Xenopus* is an aquatic species, the recorded heat temperature variations are likely overestimated, as water acts as a thermal buffer. Indeed, stage 43/44 tadpoles did not survive at 37 °C but remained viable and active at 32 °C (Table 1). Therefore, we analyzed the skin pigmentation response to temperature fluctuations between 16 °C and 32 °C.

**Table 1:** Temperature selection based on the November and December temperatures registered for three national parks located in southern Africa. † Indicates embryos do not survive at this temperature

|  |  | Cold |  | Hot |  |
| --- | --- | --- | --- | --- | --- |
|  |  | Mean daily min. | Cold nights | Mean daily max. | Hot days |
| Etosha | Nov. | 18 | 12 | 36 | 40 |
| (Namibia) | Dec. | 20 | 14 | 35 | 39 |
| Kruger | Nov. | 19 | 14 | 30 | 38 |
| (South Africa) | Dec. | 20 | 15 | 30 | 37 |
| Hwange | Nov. | 23 | 18 | 35 | 39 |
| (Zimbabwe) | Dec | 22 | 18 | 33 | 38 |
| Average |  | 20.3 | 15.2 | 33.2 | 38.5 |
| Temperature | tested | 16 |  | 37 | † |
| Temperature | selected | 16 |  | 32 |  |

### Heat induces a fast, reversible, and systemic skin darkening

We asked if *Xenopus* tadpoles darken their skin in response to warmer temperatures (16 °C to 32 °C). This paradigm is opposite to that previously used in our laboratory, where cooling conditions (24 °C to 6 °C) induced skin lightening mediated by Trpm8 (Malik et al., 2023). Importantly, we took two approaches to ensure temperature was the only variable that changed in our experiments. First, the analysis of pigmentation variation by temperature was always performed at the approximate middle of the light phase (Zeitgeber +5 to +7) since skin color changes in a daily circadian manner (Bertolesi et al., 2025). Second, the color of the surface (white) and the overhead illumination (approximately 1000 lux) were always the same to avoid pigmentation changes induced by background adaptation (Bertolesi et al., 2020, 2022). We observed that switching tadpoles from 16 °C to 32 °C induced rapid skin darkening that reached a maximum response after 40-45 minutes and resulted in an increased pigmentation index (Fig. 1 A). This response was reversible, with lightening occurring when tadpoles were moved back to 16°C (Fig. 1 A; Rev). The darkening in warm conditions and the reversibility was true for melanophores located in different body regions, including the head, belly, and tail (Fig. 1 B and 1 C), suggesting a systemic response. Finally, we observed that the pigmentation index adjusted to the environmental temperature. When we moved tadpoles gradually from 16 °C to 32 °C over a one-hour period and measured the head pigmentation index, gradual skin darkening was detected in a temperature-dependent manner (Fig. 1 D). Of note, the smallest temperature change that was needed to significantly impact skin color varied between hatches; We detected a significant response as low as 22 °C (in 1 out of 3 experiments), with the response continuing to progress until the maximum tested temperature (32 °C) (Fig. 1 D).

**Figure 1:**
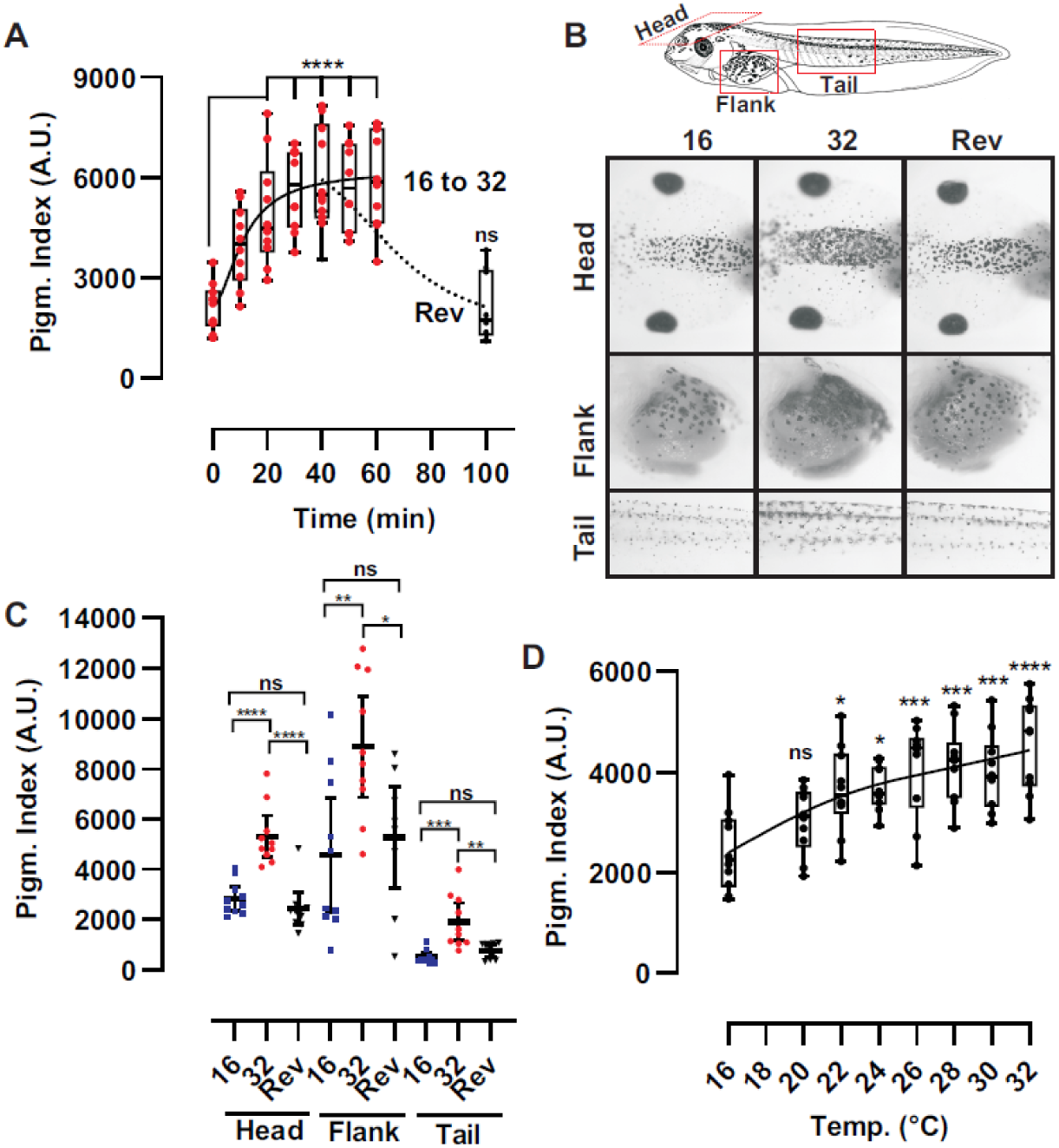
Darkening of skin colour triggered by warm temperatures is temperature dependent, fast, reversible, and a systemic response of skin melanophores. **A**) Time dependent change in pigmentation index quantified from digital images of the dorsal head of *Xenopus* stage 43/44 tadpoles moved from 16 °C to 32 °C degrees; [individual data points (n=10 embryos) and a box plot (25^th^ to 75^th^ percentile) are represented; A representative study from three (N=3) independent experiments is shown]. **B, C**) Schematic of stage 42/43 tadpole and representative pictures (**B**) and pigmentation index (**C**) from the dorsal head, lateral belly (flank) and the tail of tadpoles at 16 °C (16; blue dots), moved to 32 °C for 45 minutes (32; red dots), or switched back to 16 °C after being for 45 min at 32 °C (Reversibility; Rev; black dots); [Horizontal bar represented the mean ± 95% Confidence Interval; n=9 embryos; A representative of three (N=3) independent experiments is shown)]. **D**) Temperature-dependent increase in the head pigmentation index of tadpoles moved from 16 °C to the indicated temperatures. [individual data points (n=8/9 embryos) and a box plot (25^th^ to 75^th^ percentile) are represented; A representative study of three (N=3) independent experiments is shown].

Together, these results demonstrate that warm conditions induce a rapid, temperature-dependent, and systemic darkening of the skin, which is reversible when the temperature is reduced.

### Cultured melanophores disperse melanosomes when the temperature rises

The systemic response observed *in vivo* led us to hypothesize a cell-autonomous response where the melanophores themselves were the sensors of high temperatures. To test this hypothesis, we switched to a melanophore cell line obtained from *X. laevis* embryos (Kashina et al., 2004) and analyzed their melanosome aggregation/dispersion in response to heat. To mimic the *in vivo* condition, the *in vitro* system was set up in 35 mm dishes with growth medium without phenol red [mimicking the colourless Modified Marc’s Ringer (MMR) solution used *in vivo*], with identical light conditions, since aggregation and dispersion of melanophores *in vitro* are affected by light (Bertolesi et al., 2025; Daniolos et al., 1990). As we showed previously, approximately half the population of MEX cells grown with 5% fetal calf serum-supplemented media exhibited melanosome dispersion (Fig. 2 A, 16 °C; white arrows), with the remaining cells showing aggregated melanosomes (Fig. 2 A, 16 °C; red arrows). Exposing the cells to a higher temperature of 32 °C induced a time- and temperature-dependent dispersion of melanosomes that reached a maximum after approximately 45 minutes (Fig. 2 A, B, and C), and which was reversible (Fig. 2 B). These results show that the melanosome dispersion induced by heat is mediated via a cell-autonomous mechanism.

**Figure 2:**
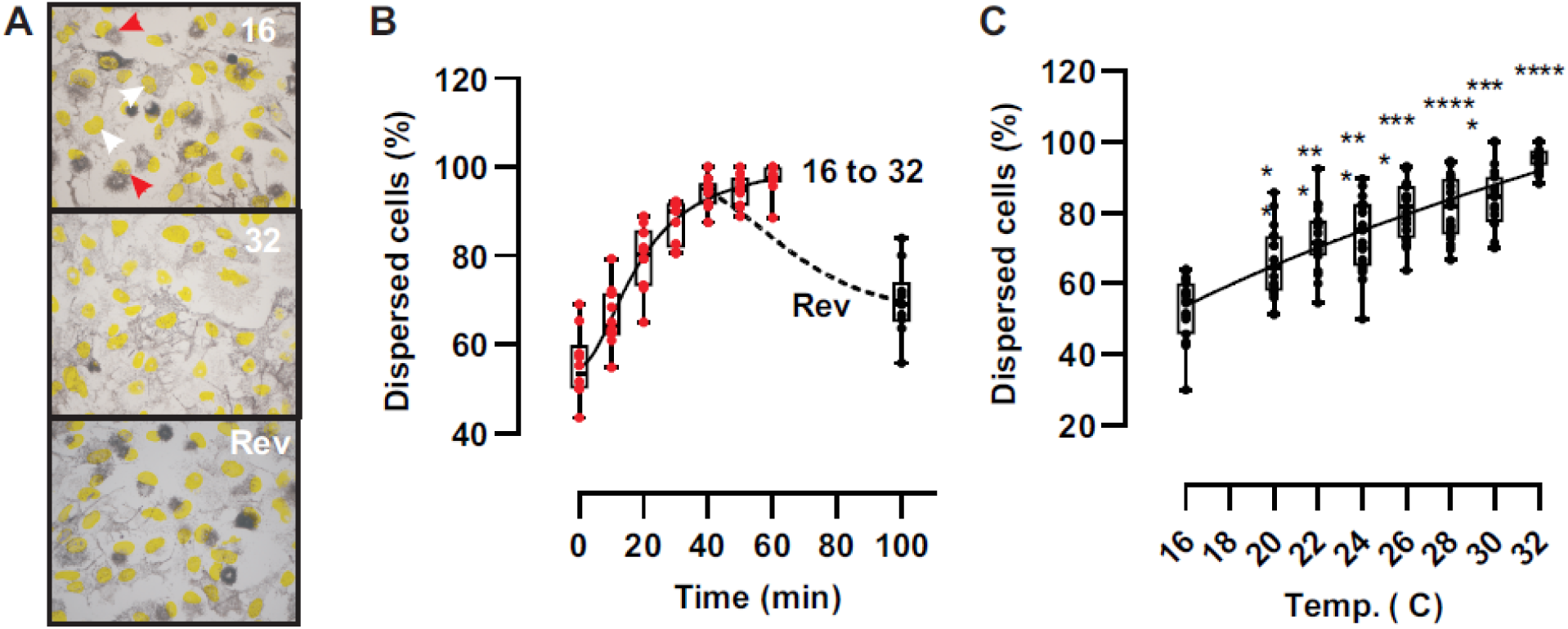
Warm temperature induces melanosome dispersion in melanophores *in vitro*. **A**) Representative brightfield images of MEX cell melanophores at 16 °C with approximately 50% of the cells showing melanosomes dispersion (white arrows) or aggregation (red arrows) at 16 °C or after 1 hour at 32 °C. Cells returned to 16 °C (Reversibility; Rev) aggregate melanosomes. DAPI stained nuclei are yellow. **B, C)** Time (**B**) and temperature dependent (**C**) increase of melanosome dispersion of MEX cells moved from 16 °C to 32 °C degrees [individual data points (n=6 pictures) and a box plot (25^th^ to 75^th^ percentile) are represented; A representative study from three (N=3) independent experiments is shown].

### Identification and expression of *trpv* and *trpa* family members in *Xenopus laevis* melanophores

To test the idea that the rapid change in skin pigmentation induced by heat in amphibians is mediated by a Trp channel(s), we first identified all *trpv* and *trpa* family members in the *Xenopus laevis* genome and analyzed their expression in melanophores. *X. laevis* is an allotetraploid species that arose via the interspecific hybridization of diploid progenitors around 18 million years ago (Session et al., 2016). Consequently, its duplicated genome contains 36 chromosomes, consisting of 18 long (L) and 18 short (S) homologous chromosomes (Session et al., 2016). While several of the duplicated genes have remained distinct, others were reduced to a single copy due to pseudogenization.

We found that most of the genes from the *trpv* family remained duplicated, except for *trpv5* and *trpv6*, each with only one copy located on the same chromosome (7L) (Fig. 3 A). Interestingly, two additional genes with unknown functions are found in the *X. laevis* genome, named *trpv4-like 1* and *trpv4-like 2*, which show high homology at the amino acid level to the predicted Trpv4 proteins (Fig. 3 A). Previous structural and molecular analysis of the Trpv4 amino acid sequence from the western clawed frog suggests that Trpv4 retained its original function, while the Trpv4-like versions diversified, with slightly different properties (Saito & Shingai, 2006). Nevertheless, these *trpv4-like* genes likely originated from intrachromosomal duplication before the generation of the *X. laevis* species, as both are present in the *X. tropicalis* genome and located on the same chromosome (6L). Finally, the *trpa1* genes were also located on chromosome 6 and remained duplicated in the *X. laevis* genome (Fig. 3 A).

**Fig 3.**
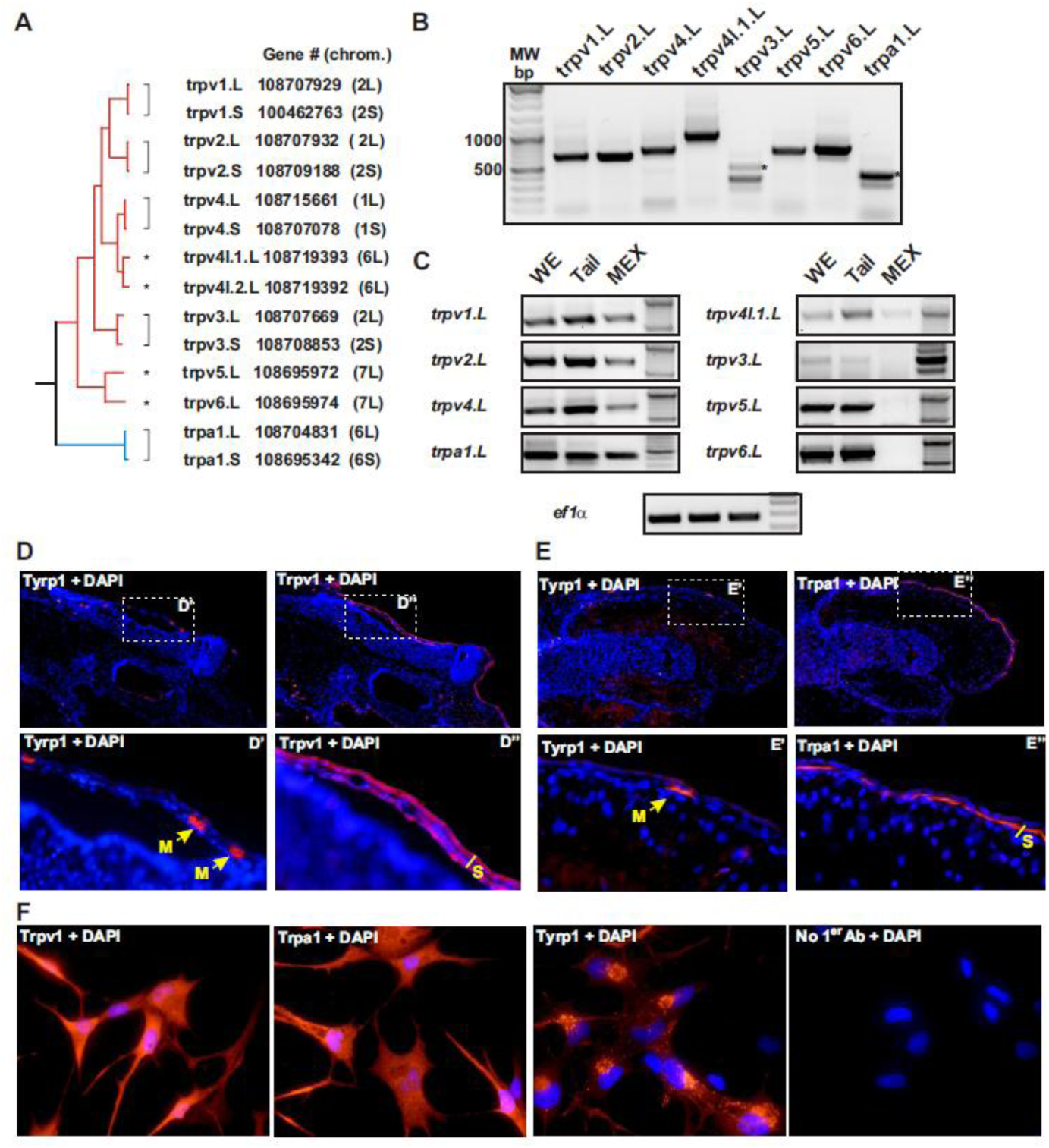
Identification and expression of *trpv* and *trpa* family members in *Xenopus laevis* melanophores. **A**) Molecular phylogenetic analysis by the maximum-likelihood method of Trpv and Trpa family members (not to scale) of the predicted proteins from the genes found in the *X. laevis* genome. The name, gene number and chromosome localization are indicated. Brackets denote genes duplicated and maintained on the long (L) and short (S) chromosomes, with single copy genes indicated with an asterisk. **B, C)** Representative RT-PCR analysis of mRNA expression in stage 43/44 whole embryos (B), and for isolated tails and MEX cells (C). The expressed mRNAs (L or S) identified after sequencing are indicated. Representative of 3 independent samples; N=3. **D-F)** Immunohistochemistry for Trpv1 (D, F) or Trpa1 (E, F) and the Tyrosinase related protein 1 (Tyrp1) in consecutive sections (D, E), and in MEX cells (F). DAPI staining (blue) was used to visualize cell nuclei and facilitate overlapping of adjacent sections in D and E. Boxed areas are shown enlarged (D’ or E’ and D” or E’’). Immunolabel indicates co-localization in melanophores (M) of the skin (S) of Tyrp1 (arrows) with the Trp channels. Note, adjacent sections were used in whole tadpoles as all antibodies were generated in rabbit. Note that Tyrp1 is localized to melanosomes while the Trp channels are in the plasma membrane.

Next, we determined which *trpa1* and *trpv* family members were expressed in whole stage 43/44 tadpoles by RT-PCR. While primers were designed to amplify both chromosomal variants (L and S) (Supplementary Table 1), cloning and sequencing of the corresponding amplicons showed exclusive expression of genes located on the L chromosomes (Fig. 3 B). To assess if melanophores express *trpa1* and *trpv* mRNAs, we also analyzed expression in isolated tails, as an *in vivo* proxy for skin, and in MEX cells. All the mRNAs were expressed in the tails, while MEX cells expressed *trpv1*, *trpv2*, *trpv4*, and *trpa1*, but little or no *trpv4-like 1*, *trpv3*, *trpv5*, and *trpv6* mRNA (Fig. 3 C). Finally, we determined for Trpv1 and Trpa1, where antibodies were available, whether the proteins were expressed by skin melanophores. We performed on tadpole frozen sections immunohistochemistry for either Trpv1 or Trpa1, with adjacent sections immunostained for Tyrosinase-related protein 1 (Tyrp1) to identify melanophores. Adjacent sections were analyzed since all antibodies were produced in rabbit. Both Trpv1 and Trpa1 were detected throughout the epidermis, including in Tyrp1-positive melanophores (Fig. 3 D and E). In support, Trpv1 and Trpa1 were also expressed by MEX cells (Fig. 3 F).

### Trpa1 is the heat sensor for melanophore dispersion with warm temperatures

To identify the TRP channel responsible for pigment dispersion by heat, we focused on the Trp channels expressed in both the tails and MEX cells: *trpv1*, *trpv2*, *trpv4*, and *trpa1*. Each Trp channel has a distinct range of thermal sensitivity and activation by chemicals (Hoffstaetter et al., 2018; Kadowaki, 2015; Zhang et al., 2022), therefore we initially performed a pharmacological study. We excluded Trpv2 from our analysis because the literature argues against this channel playing a role in thermosensation at environmentally relevant temperatures; In mammals, TRPV2 is only activated by extreme heat (52°C) and *Trpv2* deficient mice display normal thermosensation (Park et al., 2011). Further, Trpv2 lacks specific agonists, with the most commonly used TRPV2 agonist, 2-Aminoethoxydiphenyl borate (2-APB), also modulating other TRP channels (Clapham et al., 2001). Thus, we analyzed the effect of specific agonists and antagonists of Trpv1, Trpv4 and Trpa1 on melanosome dispersion of tadpoles *in vivo* and MEX cells *in vitro*.

Piperine, the Trpv1 active compound found in black pepper, did not darken the skin of tadpoles when added at 16 °C (Fig. 4 A). Piperine also did not change melanosome distribution in MEX cells (Fig. 4 A), suggesting Trpv1 is not involved in skin darkening. TRPV4 is expressed in mammalian melanocytes (Zheng et al., 2019), yet neither the pigmentation index *in vivo* nor the dispersion of melanosomes *in vitro* were significantly affected by GSK1016790A, a potent and selective TRPV4 agonist (Fig. 4 A). Additionally, Trpv1 (capsazepine) and Trpv4 (GSK2193874) antagonists failed to block the melanosome dispersion induced by switching tadpoles and MEX cells from 16 to 32 °C, at doses that were not toxic to the embryo (Fig. 4 B). These results suggest that Trpv1 and Trpv4 are not the thermosensors involved in melanosome dispersion.

**Figure 4:**
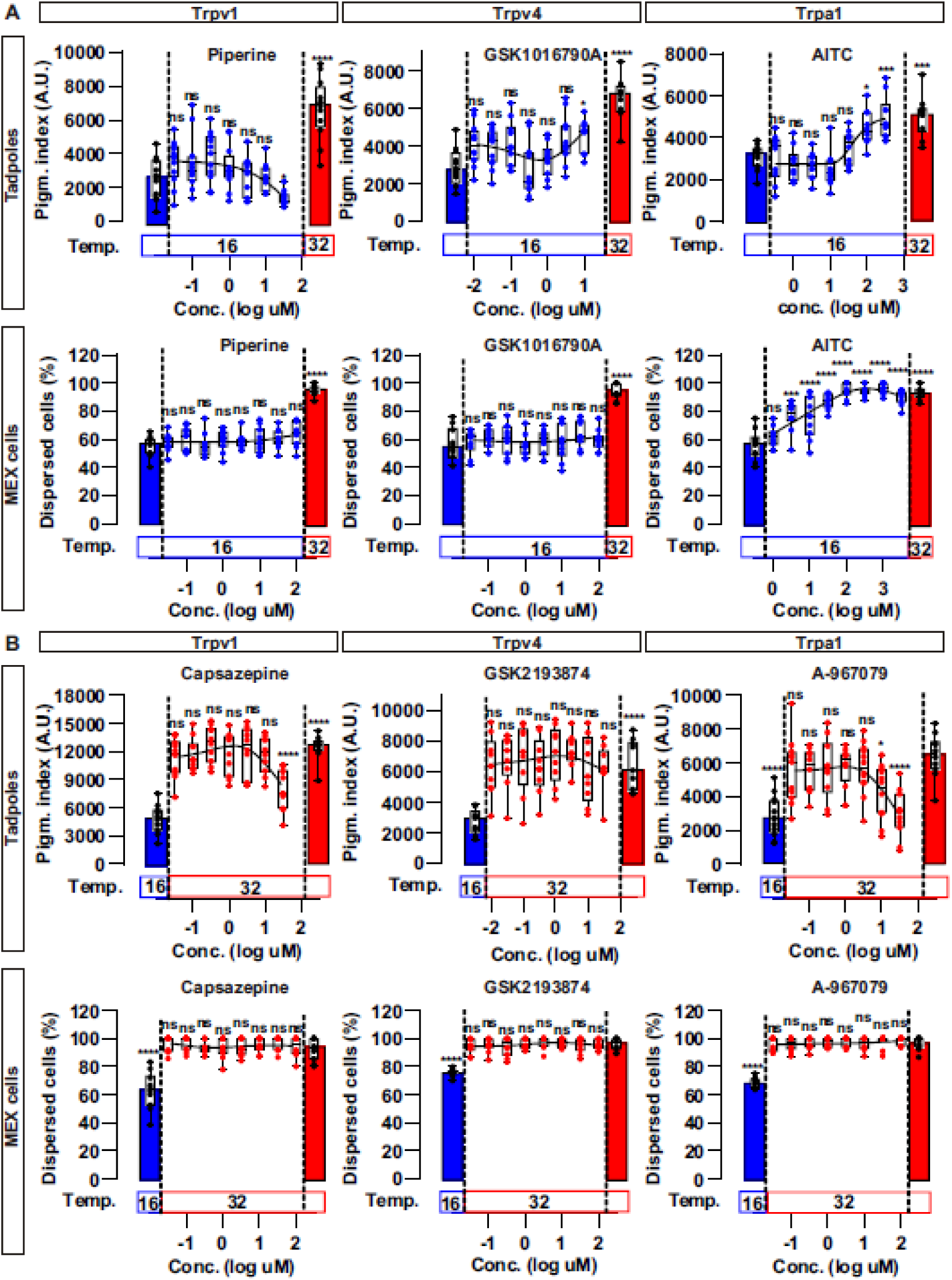
Melanosome dispersion is affected by a Trpa1 agonist and antagonist. Tadpoles (Stage 43/44) or MEX cells were treated with agonists (**A**) or antagonists (**B**) against the indicated Trp channels. The agonists were added at 16 °C and maintained at that temperature while antagonists were added at 16 °C and embryos then switched to 32°C for 45 minutes. Statistical significances for the different compounds are indicated against the control without the drug at the same temperature. ns; nonsignificant. *; p<0.05. **; p<0.01.***; p<0.001. ****; p<0.0001.

Although the thermal sensitivity of TRPA1 differs among distantly related animal species, chemical sensitivity to electrophilic compounds, particularly the Trpa1 agonist, Allyl isothiocyanate (AITC; oil of mustard), is conserved; demonstrated through electrophysiological studies in zebrafish (Oda et al., 2016), frogs (Saito et al., 2012), reptiles (Gracheva et al., 2010; Saito et al., 2012), birds (Saito et al., 2014), rodents, and humans (Chen et al., 2013). Interestingly, AITC induced a dose-dependent dispersion of melanosomes *in vivo* and *in vitro* (Fig. 4 A), suggesting Trpa1 activation induces a similar dispersion response to that seen with heat. To confirm a role for Trpa1, we next asked if a Trpa1 antagonist blocked the melanosome dispersion induced by switching tadpoles and MEX cells from 16 to 32 °C. Identifying a useful TRPA1 antagonist is difficult, in that the known antagonists show striking species-specific differences. For example, A-967079, a potent mammalian antagonist (Paulsen et al., 2015), acts as an agonist in chicken and has no effect in *X. tropicalis* heterologous overexpression systems (Banzawa et al., 2014; Nakatsuka et al., 2013; Saito & Tominaga, 2017). Interestingly, we observed a partial antagonist response in *X. laevis* tadpoles (Fig. 4 B), but were uncertain if this did not reflect toxicity (Fig. 4 B). *In vitro*, however, A-967079, showed no effect on melanosome dispersion of MEX cells (Fig. 4 B).

Thus, to confirm that Trpa1 activation causes melanosome dispersion, we turned instead to siRNA-mediated knockdown of Trpa1 in MEX cells. The expectation would be that *trpa1* siRNA should block the dispersion response to heat. Preliminary assays on MEX cells showed a transfection efficiency of less than 20% when using GFP expression vectors, making it difficult to determine the siRNA efficacy on the entire MEX cell population in culture. Therefore, we co-transfected siRNA with GFP at a 5:1 ratio using micelles of lipofectamine 2000 and analyzed only the heat response of GFP-expressing cells. GFP+ melanophores expressing sense-siRNA (S) and cells transfected only with a GFP control plasmid showed normal melanosome dispersion in response to heat (Fig. 5 A and 5 B). In contrast, GFP+ melanophores failed to disperse their melanophores when switched from 16 to 32 ℃ when Trpa1 was silenced with antisense-siRNA (AS). These results argue strongly that Trpa1 activation in *Xenopus* ectotherms is triggered by temperature and induces melanophore darkening.

**Figure 5:**
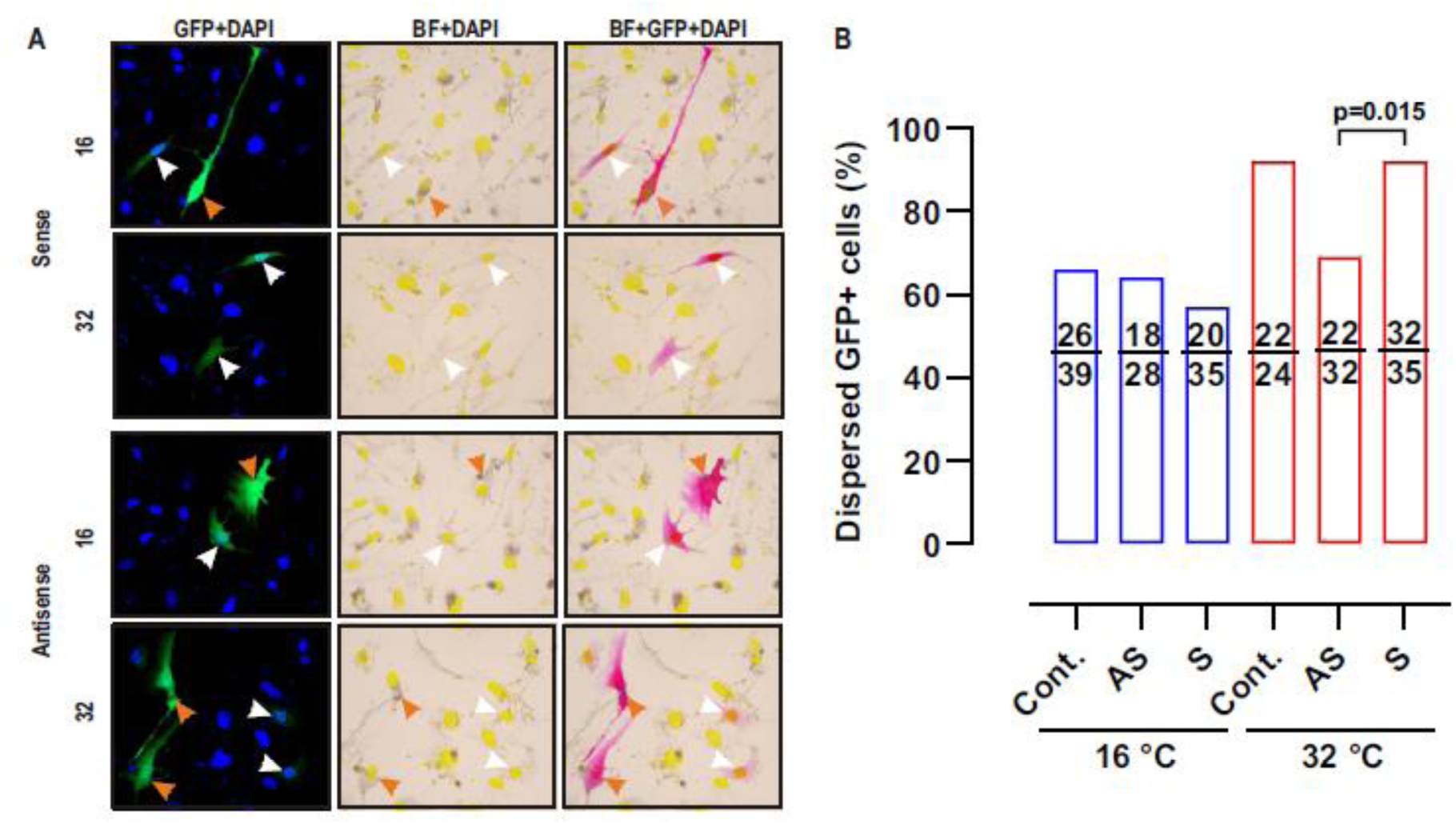
Melanosome dispersion induced by temperature is mediated by Trpa1. **A**) MEX cells transfected with a GFP-expressing vector (control) or siRNA oligonucleotides, sense (S) or antisense (AS) against Trpa1, together with GFP in a 5:1 ratio. GFP-expressing cells had aggregated (orange arrow) or dispersed (white arrow) pigment. Representative images show the merge between GFP (pink), DAPI (yellow), and brightfield (BF). **B)** Quantification of melanosome dispersion in GFP expressing cells observed by shifting cells from 16 ℃ to 32 ℃ for 1 hour after 48 hours of transfection. Data are expressed as the percentage of GFP positive cells with dispersed melanosomes relative to the total number of GFP positive MEX cells (N=3; n=20 pictures). Numbers in the bars indicates the counted cells. P value indicates a Fisher exact test comparing S and AS at 32 ℃.

### Melanogenesis and Trpa1 thermal sensitivity are linked to the evolution of thermoregulation

These Trpa1 data indicate for the first time in an ectotherm that Trpa1 functions in skin pigmentation responses to heat. These results differ from findings in hairy rodents where the channel is particularly sensitive to mechanical and noxious chemical cutaneous stimuli rather than temperature (Zappia et al., 2017). Moreover, TRPA1 thermosensitivity to cold rather than heat was demonstrated by using TRPA1 deficient mice and rats (Kwan et al., 2006; Reese et al., 2020). Finally, while TRPA1 does promote skin pigmentation in humans, it is UV light rather than temperature that induces tanning through melanin synthesis (Bellono et al., 2013; Wu et al., 2023). Taken together these data suggest an evolutionary shift in the role of TRPA1 in skin physiology. Melanin and Trpa1 likely share an important thermoregulatory role in ectotherms (Moreno Azócar et al., 2020; Stuart-Fox et al., 2017) and extinct aquatic tetrapods that used skin pigmentation to manage their physiology for survival in cold environments (Lindgren et al., 2014, 2015; McNamara et al., 2021). In contrast, the function of TRPA1 as a thermosensor may have co-evolved with the advent of integumentary coverings such as hair and feathers for thermoregulation, and possibly in hairless hominids of the African savanna with the need for the skin to provide UV protection. If true, changes in organismal thermoregulation/thermal-sensitivity and skin physiology should be reflected in the TRPA1 molecular structure.

Thus, we compared in distinct extant species the molecular structure of TRPA1 as related to the temperature range of the channel as identified previously by electrophysiology. Of note, mechanical and chemical stimuli were not considered in our analysis. The molecular structure of TRPA1 consists of six transmembrane α-helices (S1–S6), a re-entrant pore loop between S5 and S6, and variable numbers (14-18) of amino terminal ankyrin repeat domains (ARD) (Paulsen et al., 2015; Saito & Tominaga, 2017; Zhang et al., 2022). For instance, the Trpa1 of *Xenopus* and humans contains 16 ARDs (Fig. 6A). A cluster of ARDs located at the amino terminal end of the protein are thought to be the “sensor domain” of TRPA1 (Cordero-Morales et al., 2011; Jabba et al., 2014; Saito & Tominaga, 2017; Zhang et al., 2022).

**Figure 6:**
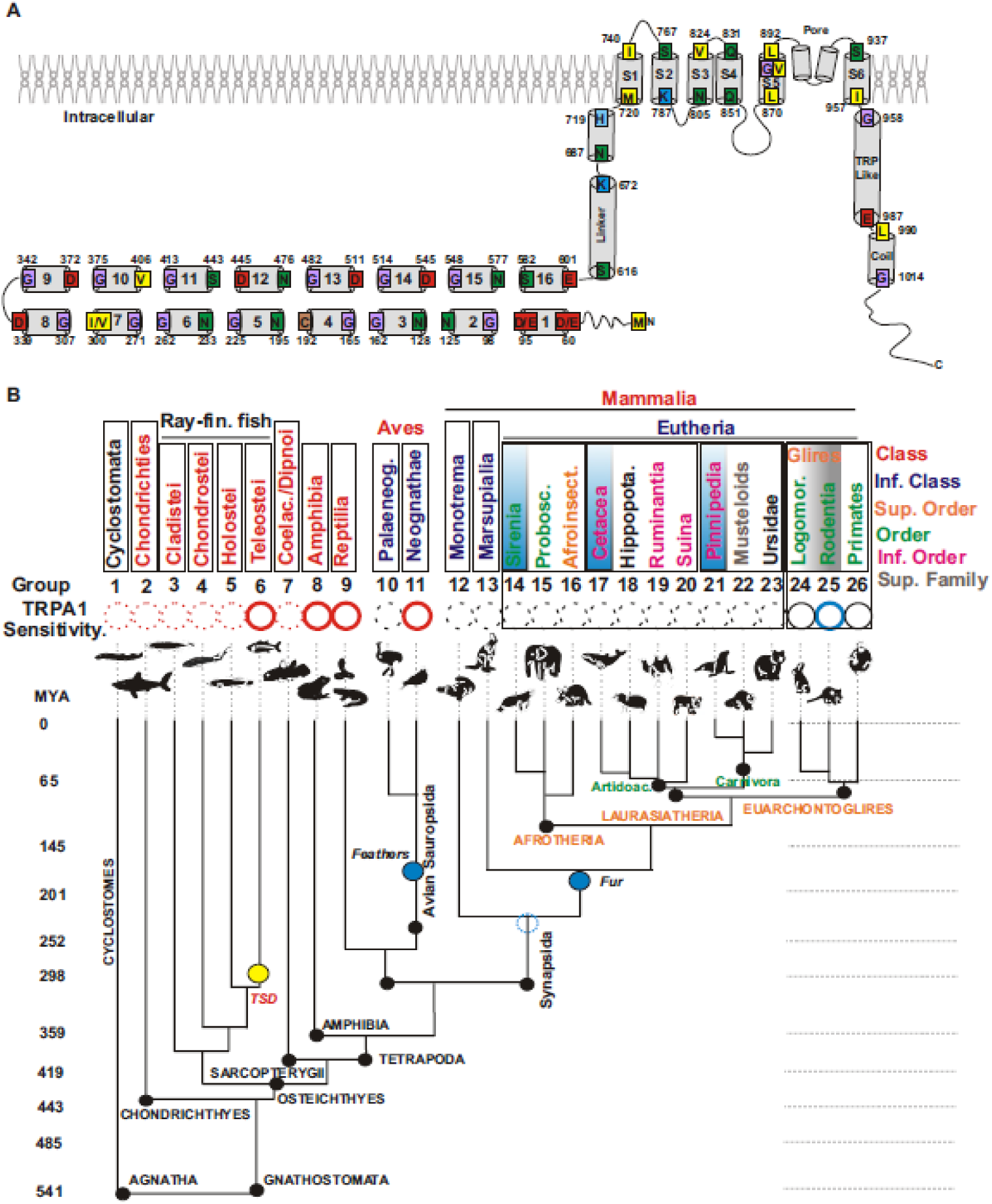
Representative TRPA1 Structure and Phylogenetic Diagram of Related Taxa. **A**) Schematic secondary structure of human TRPA1: The 16 ankyrin repeat domains (ARD) at the amino terminus are numbered, followed by the linkers, the six transmembrane domains (S1 to S6), the TRP like domain and the coiled coil domain at the carboxy terminus. **B**) Related taxa (classes or superfamily) of vertebrates with similar TRPA1 ‘thermosensation’ as determined by electrophysiological data obtained from the literature are color coded (heat, red; cold, blue; temperature insensitive, black and unknown, dashed circles). Extant species of interest shown separated (see Materials and Methods) into 26 groups corresponding to the coloured taxonomic clades indicated on the right. Evolutionary years (millions) dividing different eras are approximated and are indicated on the left. Teleost-specific duplication [TSD] is indicated in yellow. Blue circles denote the age of the oldest preserved fossils with fur or feathers with insulation capacity, likely related with the advent of homeothermy (groups 10 to 26). Group 1 to 9 are ectotherms. Groups 14 to 23 (square) are analyzed in Figure 8 and contain several specific examples of marine mammals, while groups 24 to 26 (square) are analyzed in Figure 7 and contain taxonomic groups with cold-activated TRPA1 (group 25, some rodents) and temperature-insensitive TRPA1 (group 24, Sciuromorpha and lagomorphs; group 26, primates).

To determine how changes in the thermal sensitivity of TRPA1 mapped onto evolutionary lineages we compared extant organisms grouped into 26 clades (taxonomic classes to infraorders) as referenced by the Integrated Taxonomic Information System (ITIS) (https://www.itis.gov/) (Fig. 6 B and Supplementary Table 2). These organisms were grouped into clades based on their shared evolutionary history and genetic similarities, providing a structural framework for analyzing thermal sensitivity trends. We used the electrophysiological literature to identify the thermal sensitivity range of TRPA1 in species across these clades. Our analysis focused exclusively on Chordata (organisms with an internal cartilaginous or bony skeleton) as the mechanisms to change skin color differ from those organisms in Protochordate. To the best of our knowledge, the temperature sensitivity of Trpa1 in lamprey (Cyclostomata; Group 1), sharks (Chondrichthyes; Group 2), and most ray-finned fish (Groups 3 to 5) has not yet been investigated by electrophysiology. Trpa1 electrophysiology was performed in Teleostei (Group 6), which possess two genes due to a specific genome duplication (TSD, Teleost specific duplication; Fig. 6 B). In several Teleostei organisms (zebrafish, medaka, and takifugu), at least one of the two Trpa1 homologues is activated by heat (Oda et al., 2016, 2017, 2018). Trpa1 is also heat-activated in amphibians (Group 8), reptiles (Group 9), and birds (Aves; Group 11) (Oda et al., 2019; Saito et al., 2012, 2014; Zhang et al., 2022) (Fig. 6 B). Interestingly, chickens (Group 11; Aves, Neognathae, including most flying birds) are the first lineage where TRPA1 may not serve as a heat sensor, as the activation threshold of chicken TRPA1 occurs at 39.4°C—a temperature below their typical body temperature (40–42°C) (Saito et al., 2014). There is no physiological data for TRPA1 in organisms from Group 10, which includes flightless birds (Aves, infraclass Palaeognathae) (Fig. 6 B). In the mammalian taxonomic class (Groups 12 to 26), TRPA1 electrophysiological data is available for placental organisms (Eutherian), revealing a thermal shift of TRPA1 with respect to ectotherms (Fig. 6 B), but is lacking for monotremes (Group 12) and marsupials (Group 13). Notably, the supra-order Euarchontoglires, which contains rodents and primates (Groups 24 to 26), exhibited a shift in TRPA1 thermal sensitivity to cold or insensitive. Finally, we were unable to find TRPA1 electrophysiological data from organisms of two additional Eutherian supra-orders, Laurasiatheria and Afrotheria (Fig. 6 B). We were particularly interested in these supra-orders in extant species that adapted to aquatic environments at different evolutionary times and with distinct thermoregulatory mechanisms. Thus, we generated additional groups (14 to 23) for further comparison (see below). Note that we had no electrophysiological data for these groups.

Several residues in TRPA1 are critical for channel opening (reviewed by (Laursen et al., 2015; Tominaga & Iwata, 2025; Zhang et al., 2022). One particularly important residue in mouse and rats is Glycine (G) 878, located within the S5 transmembrane domain. For human TRPA1, however, this residue is a valine (V). This amino acid difference, G878 in rodents versus V878 in humans, was suggested as an explanation for the thermal shift of TRPA1 from cold-sensitive to insensitive (Chen et al., 2013) (Fig. 6 A and Fig. 7). We compared the amino acid sequence of TRPA1 from Primates and Dermoptera (group 26) and Glires (groups 24 and 25), which refers to the common ancestry and evolutionary relationship between rodents (order Rodentia) and lagomorphs (order Lagomorpha; group 24 includes rabbits, hares, and pikas) (Fig. 6 B). V878 was present in all primates and lagomorphs, as well as in the rodent lineage that generated squirrels (suborder Sciuromorpha) (Fig. 7). A substitution at this residue (878 V to G) occurred in the rodent lineage that generated Myomorphs (mice and rats), Castorimorphs (castors) and Hystorimorphs (mole rats) (Fig. 7). Interesting, these lineages emerged after squirrels diverged during rodent evolution (Blanga-Kanfi et al., 2009). Cold temperatures activate Trpa1 in mice, rats and guinea pigs, but not in squirrels and humans (Chen et al., 2013; Matos-Cruz et al., 2017; Tominaga & Iwata, 2025; Zhang et al., 2022). Note, however, that this residue is necessary but not sufficient on its own to explain the thermal shift observed in mammals, in that an engineered conversion of V878 to G878 in human TRPA1 does not induce cold sensitivity (Chen et al., 2013).

**Figure 7:**
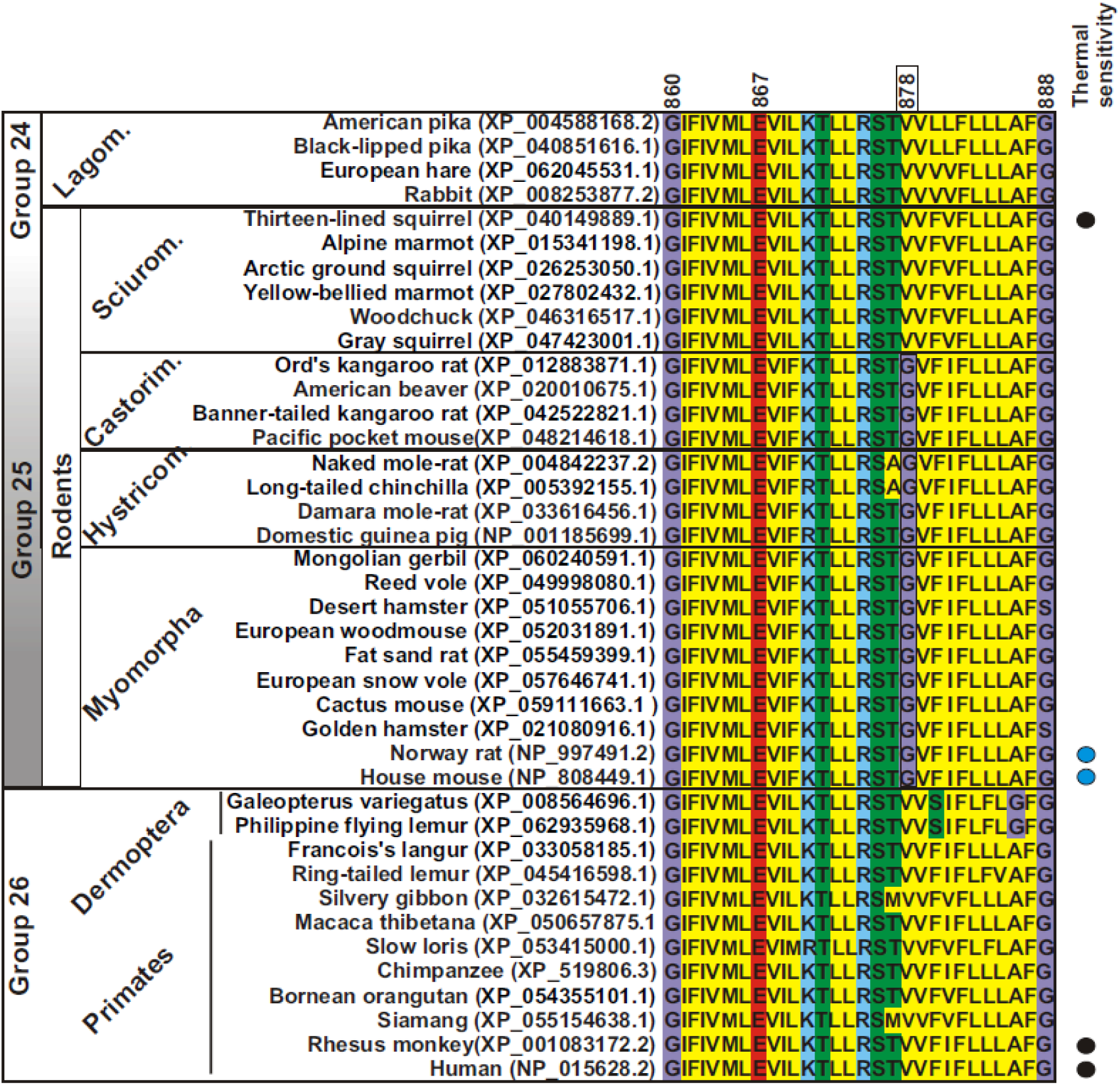
Alignment of the S5 Region in Various Species as Indicated in Figure 6. The glycine residue (G878) present in several rodents (Group 25: Myomorphas, Hystricomorphas and Castorimorphas) is essential for cold sensitivity (Chen et al., 2013). The substitution of this residue with valine (V) in primates (Group 26) and lagomorphs (Group 24: rabbits, hares, and pikas) explains the thermal differences in TRPA1 sensitivity. Substituting human V878 with G878, however, does not restore cold sensitivity. Note that TRPA1 in the rodent squirrel (Sciuromorpha) is temperature insensitive and contains V at amino acid 878 instead of G (Matos-Cruz et al., 2017). Demonstrated electrophysiological data for TRPA1 thermal sensitivity (cold, blue; insensitive, black) are indicated on the right.

While a single residue at the pore, like G878, is essential for mouse TRPA1 cold sensitivity, several studies suggest the domain responsible for thermosensing is located in the ARD region. First, the three-dimensional structure of human TRPA1 obtained by single-particle electron cryo-microscopy shows a close proximity between the ARD and helix-turn-helix motifs that allow the ARD to transmit information to the pore (Paulsen et al., 2015). Second, an unbiased random mutational analysis performed in mouse TRPA1 shows that alteration of three amino acids in ARD6 can be altered to switch thermal sensitivity from cold to heat (Jabba et al., 2014). Moreover, electrophysiology shows that two snakes with significantly different thermal activation temperatures for Trpa1 (28°C and 38°C) experience thermals shifts when the amino terminal regions of their proteins are interchanged (Cordero-Morales et al., 2011). Finally, when the thermally insensitive human TRPA1 is engineered to contain the first 10 ankyrin repeats from a rattlesnake the protein gains heat sensitivity (Cordero-Morales et al., 2011).

As proteins evolved from a common ancestor, their sequences and structures diverged (Chothia & Lesk, 1986). Importantly, the variation in mutation rate of amino acids within different domains of the same protein often depends on the functional roles of the domains. For example, in TRP channels, a “fingerprint” of mutations in the S1-S6 transmembrane domains is linked to the evolutionary selection of ion channel selectivity (Cabezas-Bratesco et al., 2022). We performed a mutational analysis for four examples of mammalian lineages that transitioned from terrestrial or freshwater aquatic environments to the sea, in that these lineages provide examples where thermoregulation evolved to rely on cardiovascular adjustments rather than skin pigmentation. In marine mammals, blood flow to the skin is often reduced during diving to conserve oxygen for vital organs, which also helps retain body heat. Conversely, increased blood flow to the skin at the surface or in warmer waters aids in heat dissipation. (Favilla et al., 2022). Additionally, the existence of a thick blubber layer for thermal insulation generally correlates with earlier aquatic adaptation, while the existence of fur is characteristic of more recently adapted organisms, as hair can increase drag in aquatic environments (Favilla et al., 2022; Khudyakov et al., 2022; Le Duc et al., 2022). We analyzed the mutation rates of TRPA1 structures associated with thermoregulatory physiology. Using the maximum-likelihood method based on the JTT (Jones-Taylor-Thornton) matrix model, which is an empirical substitution model commonly used in evolutionary studies to estimate amino acid replacement rates, we compared three regions from the full-length TRPA1 (TRPA1 full length): 1) the amino-terminal region that contains nine ARDs (NH2-ARD 1-9) and likely where thermal sensitivity resides; 2) a middle domain with ARD10-16, plus two linker domains (ARD10-16-linker); and 3) the transmembrane regions, where the pore is localized, and carboxy-end (S1-S6-COOH) regions (Fig. 6 A and Fig. 8 scheme). We hypothesized that the rate of amino acid change in the sensory region (ARD1-9; Fig. 6 A) of TRPA1 would be highest in lineages that transitioned earlier to the sea. We also predicted, in contrast, that mutation rates for the transmembrane S1-S6 domain containing the pore for cation influx would remain low in all four lineages, as this domain is linked to the evolutionarily conserved selectivity of ion influx for non-selective cation channels (Cabezas-Bratesco et al., 2022). Indeed, the mutational rates in the pore region were low (Rate close to 1) (Fig. 6 A).

**Figure 8:**
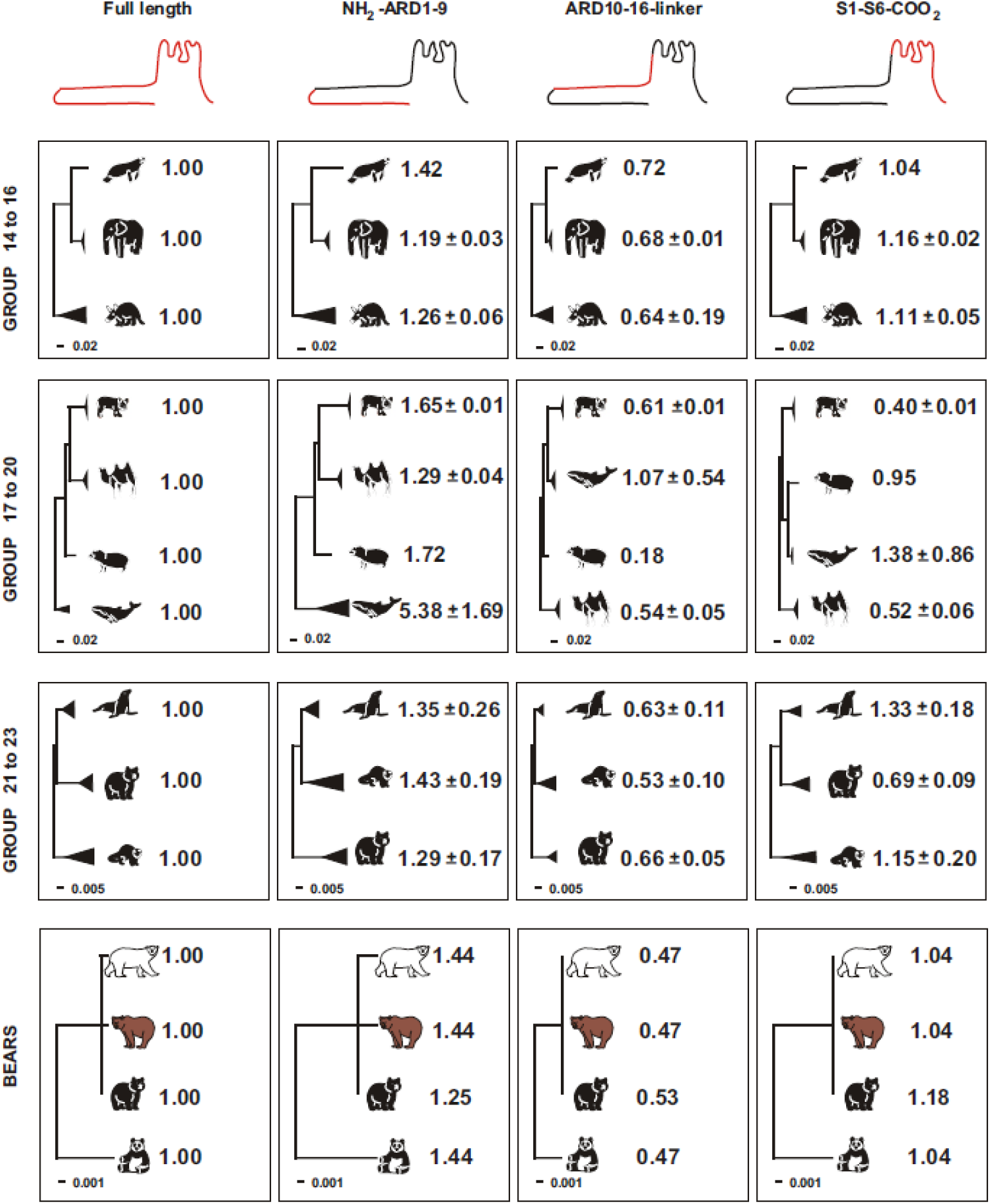
Comparative phylogenetic analysis of TRPA1 domains for three marine mammals with terrestrial relatives. Molecular phylogenetic analysis by the maximum-likelihood method based on the JTT matrix model were performed with the amino acid sequences of species shown in Supplementary table 2. The trees with the highest log-likelihood are shown in scale. Note the scale changes between the more recent evolutionary event containing bears (polar bear; brown bear; black bear and panda) and seals (Pinnipeds; groups 21-23) with respect to the analysis with manatees (Sirenia; group 14 to 16) and whales (Cetacea; group 17 to 20). The full length TRPA1 sequence, the region containing the sensor with the ankyrin repeat domain (ARD) 1 to 9, ARDs 10 to 16, and the transmembrane domains (S1 to S6) were used for the analysis. Numbers in the right (mean ± SD) indicate the phylogenetic distance relative to that for TRPA1 full length. Lack of standard deviation indicates only one species with TRPA1 data available. Evolutionary analyses were conducted in MEGA X (Kumar et al., 2018).

From the numerous independent evolutionary events in which mammals adapted to aquatic environments, we focused on four groups of extant species that exhibit distinct skin characteristics: 1) Sirenians (manatees and dugongs), also known as sea cows, possess thick, wrinkled skin covered with short, fine sensory hairs. Their blubber layer is notably thinner as compared to other marine mammals, likely due to the warmer aquatic niche that they inhabit (20–30 °C) (Le Duc et al., 2022). Having adapted to an aquatic environment approximately 50 million years ago (MYA), manatees share an evolutionary relationship with two clades of extant terrestrial species: elephants (order *Proboscidea*) and Afroinsectiphilia (African insectivores), which include golden moles, aardvarks, and Cape elephant shrews [(Groups 14–16) (Fig. 6B, Fig. 8, and Supplementary Table 2)]. 2) Cetaceans (whales, dolphins, and porpoises) which are entirely adapted to a cold aquatic lifestyle, with a smooth skin (hairless) and a thick layer of underlying blubber. This blubber provides insulation, buoyancy, and energy storage and is highly vascularized (Espregueira Themudo et al., 2020). Cetaceans became aquatic from a common terrestrial ancestor of hippos and whales (Whipomorpha) at approximately the same evolutionary time as Sirenians (50 MYA), diverging from a lineage that includes extant terrestrial species such as peccaries (Suina) and ruminants (Ruminantia) [(Group 17 to 20) (Fig. 6 B, Fig. 8, and Supplementary Table 2)]; 3) Pinnipeds (seals, sea lions, and walruses), like Cetaceans, have a skin with a thick blubber layer, but differ in that their integument is covered with short, dense fur, likely reflecting a recent adaptation to an aquatic environment (20 MYA) (Khamas et al., 2012; Khudyakov et al., 2022). Pinnipeds can additionally regulate body temperature through spending time on land, mainly during molting (Gentry, 1973; Thometz et al., 2023). Pinnipeds are carnivores that are evolutionarily related to minks, otters, and ferrets (Musteloidea), and bears (Ursidae) [(Group 21 to 23) (Fig. 6 B, Fig 8, and Supplementary Table 2)]. Finally, 4) Polar bears, considered marine mammals due to their reliance on the marine environment for hunting. Polar bears lack a typical blubber layer, possessing a dense fat layer in the skin with a distinctive composition, alongside the ability to secrete fur sebum with anti-icing properties (Carolan et al., 2025). Additionally, polar bears and sea otters are insulated by air trapped in their thick fur, which is covered with millions of hairs per square inch (Leroy et al., 2024; Strobel et al., 2022). We compared TRPA1 from polar bears with their terrestrial bear relatives (Brown bear, black bear and giant panda) [(Group 23). (Fig. 6 B, Fig. 8, and Supplementary Table 2)].

We analyzed the phylogenetic distance between these four groups and compared the full-length TRPA1 protein against the three regions of interest (expressed as a ratio: region of interest / full length) across representative members of each group. For all four groups, the aquatic and terrestrial members of the group appeared to show similar low mutation rates in the S1-S6 and ARD10-16 linkers (Fig 8). In contrast, those lineages that became aquatic early in evolution, the manatees and whales, appeared to show higher mutation rates in the amino acid sequence of their ARD1-9 in comparison to their terrestrial relatives [ratios: Group 14 (1.42; manatees) vs. Group 15 (1.19; elephants); Group 17 (5.38; whales) vs. Group 18 (1.72; Hippo); Fig. 8)]. In contrast, for seals/sea lions and polar bears, which evolved more recently, the mutational rates in the ARD1-9 region were not higher, when expressed as a ratio to the mutational rates of the full-length TRPA1, to those obtained for their corresponding relatives [ratio: Group 21 (1.35; seals/sea lions) vs. Group 22 (1.43; Musteloidea) and 1.44 for polar, panda and brown bears)].

Together, these results demonstrate that the thermosensitivity of TRPA1 has changed over the course of evolution. The protein shifted from functioning as a heat sensor in fish, amphibians, and reptiles to likely losing its noxious heat function in birds, and eventually becoming either thermally insensitive or a cold thermosensor in mammals. This shift correlates with the emergence of thermoregulation and the development of feathers and fur in the integument of homeotherms. In rodents (e.g., mice and rats), TRPA1 functions as a cold sensor. However, TRPA1 in the squirrel lineage (also rodents) and primates appears to be thermally insensitive. Finally, we observed an increased mutational rate in the sensory region of TRPA1 in early adapted marine mammals (e.g., manatees and whales) compared to their terrestrial relatives.

## DISCUSSION

Changing skin color is a crucial physiological process that aids in the thermoregulation of ectotherms. We demonstrate that TRPA1 functions as a heat sensor, regulating temperature-induced changes in skin pigmentation in *Xenopus laevis*. These data provide strong support for the idea that TRPA1 thermal sensitivity correlates with integumentary traits. TRPA1 serves as a heat sensor in ectotherms with uncovered integuments (e.g., fish, amphibians, and reptiles), and as a non-noxious thermal sensor in feather-covered birds. We propose that TRPA1 became thermally insensitive in the Euarchontoglires lineage and subsequently evolved cold sensitivity in several lineages of rodent. Our analysis suggests that TRPA1 of marine mammals experienced reduced selection pressure in lineages that adapted early (e.g., manatees and whales) with thick, sparsely haired skin, but that organisms covered with dense hair who adapted more recently to marine environments (e.g., sea lions and polar bears) show lower mutation rates in the TRPA1 thermosensor region. Skin is the largest ‘organ’ of an organism’s body with essential thermoregulatory functions. We propose that TRPA1 function in skin physiology and its thermal sensitivity evolved hand in hand during evolution.

### Trpa1 regulates melanosome dispersion in amphibians, impacting pigmentation as a thermoregulatory mechanism

In this study, we observed that *Xenopus laevis* tadpoles darken their skin rapidly in response to moving to an environment with an ecologically relevant warm temperature (32°C). This darkening occurs via melanosome dispersion within melanophores, and is both reversible and regulated in a cell autonomous manner. We propose that Trpa1 acts in melanophores to mediate the dispersion of melanosomes. In support, we find mRNA for Trpa1 in tadpole tails and melanophore MEX cells. Further, Trpa1 is co-expressed with Trpv1 by skin melanophores. While melanophores co-express Trpa1 and Trpv family members, pharmacological studies with specific agonists and antagonists, combined with molecular knockdown of Trpa1, argue strongly that Trpa1 induces the skin darkening response to heat. In support of Trpv1 playing no role, the activation threshold for *X. laevis* Trpv1 is approximately 40°C, and calcium influx through the channel is undetectable at 34°C for *X.laevis* Trpv1-transfected HeLa cells (Saito et al., 2016), both temperatures higher than our thermal-testing paradigm.

In *X. laevis*, we find two *trpa1* genes: *trpa1.L* and *trpa1.S*, though we find only the long form appears to be expressed in *Xenopus* melanophores, as suggested by previous work showing that while *trpa1.L* is expressed in the brain, spinal cord, skin, and peripheral neurons, *trpa1.S* mRNA is restricted to the brain (Saito et al., 2016). *Xenopus* Trpa1 appears sensitive to temperatures in a complementary range to that of Trpm8, with the former mediating skin darkening in response to heat, and the latter skin lightening in response to cold conditions (Malik et al., 2023). Interestingly, activation of Trpm8 also impacts tadpole locomotor performance, while anuran tadpoles across several species exhibit Trpa1-mediated heat avoidance behavior (Saito et al., 2022). Thus, pigmentation and thermal behavioral responses appear connected, with hot and cold temperatures sensed by Trpa1 and Trpm8, respectively. Note that the temperature paradigm we chose was based on the thermal characteristics of the *Xenopus laevis* niche. Although Trpa1 functions as a heat sensor in ectotherms, small differences in Trpa1 thermal sensitivity exist even between closely related species. For example, *Xenopus tropicalis*, found in western tropical Africa, has a higher Trpa1 activation threshold than *Xenopus laevis*, which is found in colder southern Africa (Furman et al., 2015; Kashiwagi et al., 2010; Saito et al., 2016). Presumably, these differences reflect distinct needs for thermal behaviour responses and/or thermoregulatory skin pigmentation in different environmental niches.

### The evolutionary shift in TRPA1 thermal sensitivity is closely linked to the thermoregulatory role of melanin in pigmentary physiology

TRP channel expression is highly conserved between ectothermic melanophores and mammalian melanocytes. Indeed, our results show that *Xenopus* melanophores express multiple and similar TRP channels, including Trpm8 (Malik et al., 2023), Trpa1, Trpv1, Trpv2, and Trpv4, to those detected in mammalian melanocytes and melanoma cells (Guo et al., 2012; Zheng et al., 2019). The conservation of Trpa1 expression between melanophores and melanocytes, alongside melanin’s role in ectothermic thermoregulation (Cuthill et al., 2017; Lindgren et al., 2014; Stuart-Fox et al., 2017) and evolutionary changes in TRPA1 thermosensitivity (Hoffstaetter et al., 2018; Laursen et al., 2015; Saito & Tominaga, 2017), suggest a connection between the emergence of endothermy and pigmentary physiology. Our systematic analysis of TRPA1 thermosensitivity based on electrophysiological data from the literature underlines the function of Trpa1 as a heat sensor in ectotherms and its modification or shift in endotherms. While we focused on Chordata, invertebrate data argues that Trpa1 originally acted as a heat sensor [Reviewed by (Hoffstaetter et al., 2018; Tominaga & Iwata, 2025)]. Our findings suggest that TRPA1 thermal sensitivity evolved alongside the advent of endothermy, correlating with changes in the localization and function of pigmented cells and melanin in skin physiology. Indeed, alterations in the location of melanin pigment-containing melanosomes over evolution are associated with changes in thermoregulatory mechanisms. For instance, in amphibians, melanosomes filled with melanin are distributed to the pigmented cells of internal organs to a much greater extent than the integumental melanophores (McNamara et al., 2021; Rossi et al., 2019), though melanin’s thermoregulatory function is likely restricted to the latter cell population. Melanin-containing pigment cells became distributed equally between the skin and internal organs of reptiles (McNamara et al., 2018), potentially enhancing the integumentary role of melanin in thermoregulation as animals transitioned to land. With the evolution of endothermy, integumentary melanosomes became dominant and pigmented cells in the integument acquired a secretory role (birds and mammals) (McNamara et al., 2018), coinciding with the evolution of feathers or hair for thermoregulation. Our analysis of TRPA1 thermal sensitivity in Chordata suggests a similar switch over evolution that coincided with changes in the involvement of the integument in thermoregulation.

The timing of the shift of TRPA1 from a heat sensor to a thermally insensitive or cold sensor during evolution remains unknown. Our findings, however, do suggest that TRPA1 was already insensitive to temperature at the base of Euarchontoglires (see below). In human melanocytes, UV light rather than temperature activates TRPA1, which may be related to adaptations in hairless hominid skin. Human melanocytes increase melanin synthesis and secretion to darken the skin, with neighboring keratinocytes capturing and transferring melanin to produce color changes. Although UV-induced melanin synthesis is conserved in vertebrates, including rodents and hominids, (Bellono & Oancea, 2013; Wicks et al., 2011; Wu et al., 2023), the underlying molecular pathways downstream of activated TRPA1 have only recently been elucidated in humans. In human melanocytes, TRPA1 activation by UV light regulates calcium influx and melanosome pH to enhance tyrosinase activity and melanin production (Jia et al., 2021; Wu et al., 2023), as well as keratinocyte phagocytosis of melanin (Wang et al., 2022). Here, TRPA1 plays a critical role in UV protection. Further studies are needed to investigate TRPA1’s role in melanin synthesis, keratinocyte phagocytosis, and UV activation across non-hominid species. Such research could reveal whether this adaptation emerged during the savanna period in response to hair loss or is conserved in hairy mammals (Dávid-Barrett & Dunbar, 2016; Ruxton & Wilkinson, 2011). This information is particularly relevant in that cutaneous-specific TRPA1 knockout mice exhibit reduced sensitivity to mechanical stimuli and temperature (Zappia et al., 2017).

### Cold thermal sensitivity in certain rodent lineages is attributed to a single amino acid substitution and explained by the phylogeny of Euarchontoglires

Interestingly, our evolutionary analyses suggest that TRPA1 was thermally insensitive in early ancestors of Euarchontoglires, but that a mutation in amino acid 878 from valine to glycine was in part responsible for conferring cold sensitivity in certain rodents. The cold thermosensing of mouse and rat TRPA1 relate to this amino acid being a glycine, in that a mouse TRPA1 version where G878 is mutated to V878 is thermally insensitive (Chen et al., 2013). In squirrels, however, temperatures as low as 10 ℃ fail to activate the channel (Matos-Cruz et al., 2017). Our evolutionary analysis of more than 50 extant rodent species reveals that the squirrel lineage retains a V878. While G878 is essential for cold sensitivity in mouse and rats, this variant is not sufficient by itself to determine cold sensitivity, as human TRPA1 does not regain cold thermal properties when V878 is switched to G878 (Chen et al., 2013). Although an independent amino acid substitution in the different lineages cannot be excluded, the evolutionary relationships within Euarchontoglires strongly support the possibility that V878 was the ancestral state in this lineage. Thus, TRPA1 was likely thermally insensitive at the base of the Euarchontoglires lineage, as V878 remains in Sciuromorphs, Lagomorphs, and primates, but mutated to G878 specifically in Myomorphs (mice and rats), Hystricomorphs (mole rats), and Castorimorphs (beavers). A phylogenetic analysis of Rodentia, based on six nuclear genes from 41 rodent species, supports this hypothesis—showing that squirrels (Sciuromorphs) diverged early, after lagomorphs and primates, while all other rodent clades emerged later from a common ancestor (Blanga-Kanfi et al., 2009).

### Reduced TRPA1 selective pressure in marine mammals correlates with the loss of fur

We find a higher mutation rate in the thermosensory region of TRPA1 correlates with the adaptation of individual marine mammals to aquatic environments and the reorganization of their integumentary structure. Marine mammals underwent distinct evolutionary adaptations, ranging from morphological (e.g., streamlined body shape, paddle-like limbs for enhanced mobility) to physiological changes (e.g., echolocation, thermoregulation). This is especially true for skin physiology, where differences—such as the presence of fur or a thick blubber layer—vary depending on when certain species adapted to a marine environment. Over evolution, mammals re-entered the marine realm at least seven separate times. Five of these lineages possess extant species (Uhen, 2007). Here, we analyzed TRPA1 of four of them. The fifth lineage corresponds to marine otters that adapted to the sea environment 5 MYA. Marine otters are carnivores of the Mustelidae family (Group 22) closely related to polar bears (Group 23; Ursidae), sharing several skin structures. Therefore, we did not analyze them as an independent group.

Our data show that the mutation rate in the thermosensory region of TRPA1 is especially elevated in the two earliest adapted (50 MYA) marine mammal lineages, the Sirenians and Cetaceans. These lineages lack typical mammalian fur and instead possess a thick skin with a substantial blubber layer used for thermal regulation (Espregueira Themudo et al., 2020; Le Duc et al., 2022). Interestingly, the smooth, drag-reducing skin of cetaceans, which is keratinized, lubricated, and hairless, resulted from specific episodes of gene loss (Espregueira Themudo et al., 2020). In agreement, we detected a higher mutational rate of cetacean TRPA1, suggesting low selection pressure. In contrast, pinnipeds, which evolved around 20 MYA, feature a fur-covered integument with a thinner blubber layer than whales (Khamas et al., 2012). The blubber of cetaceans and pinnipeds is highly vascularized, in part to regulate thermal dissipation (Favilla et al., 2022). Note that pinnipeds also thermoregulate by spending time on land (Gentry, 1973; Thometz et al., 2023). Pinnipeds, as well as polar bears, show a mutation rate in the TRPA1-thermosensitive region that appears to be similar to that of their terrestrial relatives. While the integument of polar bears is highly pigmented (melanin), solar energy does not reach the skin epithelium of the thick dorsal fur, and only partially does in thinner fur regions (Leroy et al., 2024). Thus, thermoregulation in polar bears appears to rely mainly on fur characteristics (Carolan et al., 2025) and the integumentary adipocyte layer, and not melanisation. Thus, in general, our data indicate less pressure selection on TRPA1 thermosensing in early adapted marine mammals with blubber and less hair, which is not the case in more recently adapted marine lineages.

TRPA1 acts as a heat sensor in amphibians, and this thermal sensitivity plays a crucial role in melanin dispersion within the integumentary melanophores of ectotherms. In organisms with uncovered integument (ectotherms), thermoregulation is partially mediated by the dispersion and aggregation of melanosomes. Interestingly, the role of integumentary melanin evolved alongside the transition to endothermy, coinciding with shifts in TRPA1 thermal sensitivity. We propose that TRPA1 function in skin physiology and its thermal sensitivity co-evolved throughout vertebrate evolution.

## Acknowledgments

This work was supported by an operating grant from the Natural Sciences and Engineering Research Council of Canada (NSERC) to SM. We thank Ms. Carrie Hehr for excellent technical assistance.

## METHODS

### Embryos, drug treatment and warm treatment

The Animal Care and Use Committee, University of Calgary, approved procedures involving frogs and embryos (AC21-0148; signed by Dr. Derrick Rancourt). Embryos were obtained by induced egg production from chorionic gonadotrophin (Intervet Canada Ltd.) injected females and in vitro fertilization according to the standard procedures (see protocols at Xenbase (http://www.xenbase.org)). Embryos were maintained at 16 °C in Marc’s modified Ringer’s (MMR) solution (100mM NaCl, 2mM KC1, 2mM CaCl_2_,1mM MgCl_2_, 5mM HEPES pH 7.4) until stage 43/44 (approximately 1 week) and staged according to Nieuwkoop and Faber on Xenbase (http://www.xenbase.org). The embryos were reared under light cycles of 12 h ON/ 12 h OFF (light =1000 lux or approximately 1.5 × 10–4W/cm^2^) on a white background. For experiments, the embryos were set in 35 mm dishes with 4 ml of MMR in a 16 °C or 32 °C incubator for the indicated times. A non-noxious warm temperature was chosen (32 °C) based on the average temperatures recorded over the last 30 years for three national parks in southern Africa (Etosha/Namibia; Kriger/South Africa and Hawange/Zimbawe) obtained from the Meteoblue website (https://www.meteoblue.com/en/weather/historyclimate/).

For immunohistochemistry, embryos at 48 h post fertilization were treated with 0.02% 1-phenyl-2-thiourea (PTU), an inhibitor of eumelanin pigment formation (Bertolesi et al., 2015). Pharmacological studies were performed with the following TRP agonists added to the MMR rearing solution containing tadpoles or MEX cells at 16 °C to determine their effect on pigmentation: 1) Piperine, a TRPV1 agonist [Abcam; (Ab142933)]; 2) GSK1016790A, a TRPV4 agonist [Millipore Sigma (G0798)] and 3) Allyl isothiocyanate (AITC), a TRPA1 agonist (Millipore Sigma; W203408). The following antagonists were added at 16 °C to MMR solution containing tadpoles or melanophores cells in growth medium, which were then switched immediately to 32°C: 1) Capsazepine, a TRPV1 antagonist [Abcam (Ab120025)]; 2) GSK2193874, a TRPV4 antagonist (Tocris Bioscience; #5106); and 3) A-967079, a TRPA1 antagonist (Millipore Sigma; SML0085). Of note, the maximum dose tested varied depending on either solubility in aqueous solution or drug toxicity. The agonists and antagonists used are organic compounds with low solubility in aqueous solutions, therefore were initially dissolved in dimethyl sulfoxide (DMSO) before dilution (1/1000) in MMR or growth medium. For example, the highest piperine dose tested was 100 µM (diluted from a 100 mM DMSO stock solution) as its maximum solubility in water is 140 µM. Toxicity causing death of tadpoles was identified by stasis after one hour that failed to reappear upon switching to a drug-free condition. Note that in general, toxicity and death were following by skin lightening.

### Determination of pigmentation index

We quantified changes in skin pigmentation by measuring skin pigmentation indices as described previously (Bertolesi et al., 2015). Briefly, pictures of the dorsal head of tadpoles were taken using a stereoscope (Stemi SV11; Carl Zeiss Canada, Ltd., Toronto, Canada) and a camera (Zeiss; Axiocam HRC), with identical conditions of light, exposure time and diaphragm aperture. Pictures were converted to binary white/black images using NIH ImageJ (U. S. National Institutes of Health, Bethesda, MD) public domain software.

### *Xenopus laevis* melanophore (MEX) cell culture

To test the pigmentation response to temperature we employed the melanophore (MEX) cell line originally generated from stage 35 *Xenopus laevis* embryos (Kashina et al., 2004). Cells were maintained in growth medium (70% Leibovitz’s L15 medium with 25% added water and supplemented with 5% fetal bovine serum (Invitrogen) without antibiotics). To mimic the conditions to observed *in vivo*, cells were maintained in medium without phenol red to maximize light penetration. For the pigmentation heat response, cultures at 16 °C were switched to 32 °C. During the warming paradigm, light from above (1000 lux) was maintained, as described previously for tadpoles. Cells were fixed with 4% paraformaldehyde and stained with DAPI (1 µg/µl) before imaging.

### Identification and expression of *Xenopus* TRP channels

The screening and identification of *Xenopus laevis trpv* and *trpa* members was described recently (Malik et al., 2023). The expression of *trp* mRNAs in whole stage 43/44 embryos and melanophores was assessed by RT-PCR. Total RNA was obtained from whole embryos, surgical isolated tails, and MEX cells using TRIzol (Invitrogen) according to the manufacturer’s protocol. Single-strand cDNA was produced from RNA samples (5 µg) by priming with oligo(dT) primers using SuperScriptTM IV reverse transcriptase (Invitrogen) according to the manufacturer’s instructions. All PCR amplifications were carried out in a total volume of 20 µL with 1 µL of cDNA, 2 µL of primers, 7 µL of water and 10 µL of 2X PCR master mix (Thermo Scientific, IL). PCR amplifications were carried out between 40 and 45 cycles and with an annealing temperature of 55 °C. PCR products obtained from cDNA were cloned into TOPO-pCRII (Invitrogen) vectors and sequenced to confirm identity. Specific primers were designed to amplify both homologous variants (on L and S chromosomes) of the different Trp channels (Supplementary table 1). The sequence of the different *trp* channels can be obtained from GenBank-NCBI database with the gene name accession numbers provided in Fig. 3 A.

### Immunohistochemistry for Trpv1 and Trpa1

Immunohistochemistry against Trpv1 and Trpa1 was performed as described recently (Malik et al., 2023) using the following antibodies: anti-crocodile Trpv1 (rabbit polyclonal; 1/200 dilution; Thermo-Fisher, Scientific; #OST00058W) and anti human TRPA1 (rabbit polyclonal; 1/200 dilution; Novus Biologicals; NB110-4076SS). The identification of skin melanophores was performed with a rabbit polyclonal antibody that recognize the Tyrp-1 (1:200 dilution; Thermo-Fisher Scientific, IL; PA5-81909)], a specific enzyme involved in melanin synthesis. Since antibodies against Trp channels and the melanophore marker (Tyrp-1) were all generated in rabbit, the analysis of co-expression was performed in consecutive 12 µm transverse frozen cryostat sections obtained from stage 43/44 embryos. Following detection with primary antibodies, slides were treated with a secondary antibody (1:1000 dilution of Alexa Fluor 488) and DAPI (1 µg/µl) to stain nuclei.

### Grouping in clades and analysis of TRPA1 thermal sensitivity from the literature

Vertebrates were divided into 26 groups based on taxon clades [(Integrated Taxonomic Information System (ITIS) (https://www.itis.gov/)] with demonstrated evolutionary relationship. Group 1 contains species in the infraphylum agnathan (cyclostomes) while Groups 2 to 26 correspond to some, but not all extant gnathostomes. We initially grouped based on the taxonomic clade “Class”. The Group 2 contains the cartilaginous fish (sharks; taxonomic class, Chondrichthyes), while the ray-finned fish (Actinopterygii) were divided into 4 groups also representing taxonomic class; The Cladistei (Group 3, bichir), Chondrostei (Group 4; sturgeons and paddlefishes) Holostei (Group 5; gars and bowfins) and Teleostei (Group 6; the largest clade of bony fish with several examples of TRPA1 cloned and characterized). The following class groups are Coelacanths and Dipnoi (Group 7; Sarcopterygii, lung fish), the Amphibians (Group 8; frogs, salamanders, and caecilians), reptiles (Group 9; lizards, snakes, turtles and crocodiles), Aves (Group 10 and 11) and mammals (Group 12 to 26). The Avian Sauropods were divided into two groups representing the ‘Inferior class’ Paleognathae (Group 10), where most primitive and mainly flightless birds reside, and Neognathae (Group 11), which include almost all living flight bird species. The three mammalian infraclasses correspond to monotremes (Group 12; egg-laying mammals), marsupials (Group 13), and eutherians (Groups 14 to 26).

Groups 14 to 23 were generated to compare three marine mammals with their terrestrial relatives: i) In the superorder Afrotheria, the Sirenians (Group 14; manatees and dugongs) were compared with elephants (Group 15; Proboscidea) and African insectivores (Group 16; aardvarks and others), ii) The cetaceans (Group 17; whales) were compared to hippopotamuses (Group 18), ruminants (Group 19), and suids (Group 20; pigs), and iii) Seals and sea lions (Group 21; Pinnipeds), marine mammals, which were compared with other carnivores including minks, otters, and ferrets (Group 22; Musteloidea), and bears (Group 23; Ursidae). The superorder Euarchontoglires (Group 24 to 26) contains most of the species where TRPA1 is well characterized, including the lagomorphs (Group 24; rabbits, hares, and pikas), rodents (Group 25; mice and rats) and primates (Group 26; human and monkeys).

### Alignment and phylogenetic analysis

Validated sequences (NCBI numbers in Supplementary table 2) were aligned using MUSCLE (multiple sequence alignment) to build a hidden Markov model (HMM) by using a maximum-likelihood architecture construction algorithm. All phylogenetic analysis and alignments were performed by using the public domain MEGA X software (https://www.megasoftware.net) (Kumar et al., 2018).

### Microscopy

Images of embryos were taken with an Axio-Cam HRc (Carl Zeiss) on the Stemi SVII stereomicroscope (Carl Zeiss). Section images were processed for brightness and contrast with Adobe Photoshop.

### Statistics analysis and reproducibility

GraphPad Prism 10.0 software was used for statistical analysis of data and graphic preparations. Statistical analysis is ANOVA followed by Bonferroni’s test. Significance was considered at p < 0.05. Experiments were performed three independent times (N= 3) unless otherwise indicated. Since pigmentation index varies between hatches, a representative experiment is shown in each figure. The independent experiments showed similar trends. Each experimental treatment contained a minimum of 9 tadpoles (n; generally, ≥9), which are represented in the figures as individual data points. Additionally, figures show a box plot (25th to 75th percentile) of the mean and 95% confidence interval. Immunohistochemical analyses were performed at less three times (N= 3), with 4 independent tadpoles (n = 4) in each experiment. CorelDraw 10.0 was used to compile multipaneled figures.

## Notes

**Competing interests:** No competing interests declared

### Competing Interest Statement

The authors have declared no competing interest.

