## Supplementary Table 1 for "Evolutionary Adaptations of TRPA1 Thermosensitivity and Skin Thermoregulation in Vertebrates"

**Supplementary Table 1:** Primers used to clone *trpv* and *trpa* genes

| Position in L | gene bank # | Position in S | gene bank # | sequence | primer identity (%) to L or S | PCR product cloned Size / name |
|---|---|---|---|---|---|---|
| *trpv1*.L/S F1498 | XM_018246044 | F1472 | XM_041582945 | gttctgcatgccctggtagac | 95 | 660 / *trpv1*.L |
| *trpv1*.L/S R2158 | | R2132 | | tgtcagcaagcaaagccttca | 100 | |
| *trpv2*.L/S F790 | XM_041581649 | F1092 | XM_018248812 | ctgctgaaggccatgtggaat | 100 | 655 / *trpv2*.L |
| *trpv2*.L/S R1445 | | R1747 | | gggccataagtccattccgtg | 95 | |
| *trpv4*.L/S F1030 | XM_018261018 | F1048 | XM_018244027 | ggcggcaaaaatgacaccatccc | 95 | 740 / *trpv4*.L |
| *trpv4*.L/S R1768 | | R1786 | | gcatctcatggcggttctcaa | 100 | |
| *trpv4l*.1/2.L F1214 | XM_018268227 | F819 | XM_018268226 | atgttcatgctcaggcctgtg | 100 | 964 / *trpv4l*.1.L |
| *trpv4l*.1/2.L R2177 | | R1848 | | aggaccagggcaatcaccata | 100 | |
| *trpv3*.L/S F653 | XM_018245670 | F612 | XM_041583285 | attgatgtacgggctcaggga | 95 | 484 / *trpv3*.L |
| *trpv3*.L/S R1137 | | R1096 | | gagtttggccagcaggtatct | 95 | |
| *trpv5*.L F522 | XM_018224919 | | | ctggtgagactgctctgcatg | | 696 / *trpv5*.L |
| *trpv5*.L R1217 | | | | tatcaaggatccgtcgtgcct | | |
| *trpv6*.L F720 | XM_018224923 | | | gcacatcctggttctacagcc | | 744 / *trpv6*.L |
| *trpv6*.L R1463 | | | | ccgtctgtgttagtgagtcgc | | |
| *trpa1*.L/S F965 | XM_041566492 | F950 | XM_018223798 | gcattttgccgctacacaagg | 100 | 427 / *trpa1*.L |
| *trpa1*.L/S R1391 | | R1376 | | atggagaggggtacagccttc | 95 | |
