## Supplementary Table 2 for "Evolutionary Adaptations of TRPA1 Thermosensitivity and Skin Thermoregulation in Vertebrates"

**Supplementary table 2:** TRPA1 genes and protein sequence ID numbers grouped as indicated in Figure 6, Figure 7 and Figure 8

| Group | ID gene | species | | ID protein | # aa |
|---|---|---|---|---|---|
| 1 | 116945229 | Petromyzon marinus (sea lamprey) | Cyclostome | XP_032815428.1 | 1081 |
| 1 | 137437431 | Myxine glutinosa (Atlantic hagfish) | Cyclostome | XP_067987847.1 | 1180 |
| 1 | 133353334 | Lethenteron reissneri (Eastern lamprey) | Cyclostome | XP_061425467.1 | 1080 |
| 2 | 109921381 | Rhincodon typus (whale shark) | Chondrichthyes | XP_048451862.1 | 1120 |
| 2 | 103174784 | Callorhinchus milii (elephant shark) | Chondrichthyes | XP_007885526.1 | 1101 |
| 2 | 116972041 | Amblyraja radiata (thorny skate) | Chondrichthyes | XP_032875200.1 | 1142 |
| 3 | 120530287 | Polypterus senegalus (gray bichir) | Cladistia | XP_039610586.1 | 1106 |
| 3 | 114653053 | Erpetoichthys calabaricus (reedfish) | Cladistia | XP_051785308.1 | 1106 |
| 4 | | Acipenser oxyrinchus oxyrinchus (Sturgeon) | chondrostei | KAK1173831.1 | 1111 |
| 5 | 102688457 | Lepisosteus oculatus (spotted gar) | holostei | XP_015208897.1 | 1136 |
| 6 | 474351 | Danio rerio (zebrafish) | teleostei | NP_001007066.1 | 1115 |
| 6 | 474353 | Danio rerio (zebrafish) | teleostei | NP_001007067.1 | 1120 |
| 6 | 113042317 | Carassius auratus (goldfish) | teleostei | XP_026056856.1 | 1108 |
| 6' | 115537365 | Gadus morhua (Atlantic cod) | teleostei | XP_030205085.1 | 1124 |
| 6' | 101174541 | Oryzias latipes (Japanese medaka) | teleostei | XP_023806150.1 | 1118 |
| 6' | 109629596 | Paralichthys olivaceus (Japanese flounder) | teleostei | XP_019942914.1 | 1133 |
| 6'' | 115103056 | Oncorhynchus nerka (sockeye salmon) | teleostei | XP_029479513.1 | 1103 |
| 6'' | 109884121 | Oncorhynchus kisutch (coho salmon) | teleostei | XP_031652805.1 | 1125 |
| 6'' | 105019660 | Esox lucius (northern pike) | teleostei | XP_034145130.1 | 1130 |
| 7 | 102366509 | Latimeria chalumnae (coelacanth) | lungfish | XP_014342061.1 | 1130 |
| 8 | 100158526 | Xenopus tropicalis (tropical clawed frog) | Amphibia | NP_001121434.1 | 1144 |
| 8 | 108704831 | Xenopus laevis | Amphibia | XP_041422426.1 | 1142 |
| 8 | 108695342 | Xenopus laevis | Amphibia | XP_018079287.1 | 1139 |
| 8 | 120941126 | Rana temporaria (common frog) | Amphibia | XP_040210341.1 | 1132 |
| 8 | 108791762 | Nanorana parkeri | Amphibia | XP_018417833.1 | 1123 |
| 8 | 121002052 | Bufo bufo (common toad) | Amphibia | XP_040289288.1 | 1124 |
| 8 | 138497614 | Ambystoma mexicanum | Amphibia | XP_069477304.1 | 1120 |
| 8 | 115085435 | Rhinatrema bivittatum (two-lined caecilian) | Amphibia | XP_029447321.1 | 1123 |
| 8 | 117353621 | Geotrypetes seraphini | Amphibia | XP_033785675.1 | 1094 |
| 9 | 103062166 | Python bivittatus (Burmese python) | Reptilia | XP_025029411.1 | 1112 |
| 9 | 100556580 | Anolis carolinensis | Reptilia | NP_001280042.1 | 1112 |
| 9 | 110089734 | Pogona vitticeps (central bearded dragon) | Reptilia | XP_020668672.1 | 1112 |
| 9 | 120308852 | Crotalus tigris (Tiger rattlesnake) | Reptilia | XP_039201373.1 | 1137 |
| 9 | 117873573 | Trachemys scripta elegans | Reptilia | XP_034618996.1 | 1115 |
| 9 | 125632552 | Caretta caretta (Loggerhead turtle) | Reptilia | XP_048696888.1 | 1114 |
| 9 | 120399538 | Mauremys reevesii (Reeves's turtle) | Reptilia | XP_039383812.1 | 1114 |
| 9 | 102944221 | Chelonia mydas (Green sea turtle) | Reptilia | XP_037748377.1 | 1114 |
| 9 | 109315826 | Crocodylus porosus (Aus. saltwater crocodile) | Reptilia | XP_019399340.1 | 1112 |
| 9 | 102575342 | Alligator mississippiensis (American alligator) | Reptilia | XP_006277080.1 | 1114 |
| 9 | 102367987 | Alligator sinensis (Chinese alligator) | Reptilia | XP_006015374.1 | 1114 |

| | | | | | |
|---|---|---|---|---|---|
| 10 | 112992219 | Dromaius novaehollandiae (emu) | Palaeognathae | XP_025970932.1 | 1126 |
| 10 | 112961286 | Apteryx rowi (Okarito brown kiwi) LQP | Palaeognathae | XP_025912075.1 | 1060 |
| 10 | 112942842 | Nothoprocta perdicaria | Palaeognathae | XP_025889658.1 | 1127 |
| 10 | 104153561 | Struthio camelus australis | Palaeognathae | XP_009687800.1 | 1192 |
| 11 | 420180 | Gallus gallus (chicken) | Neognathae | NP_001305389.1 | 1126 |
| 11 | 128142002 | Harpia harpyja (harpy eagle) | Neognathae | XP_052643510.1 | 1126 |
| 11 | 128905636 | Rissa tridactyla (black-legged kittiwake) | Neognathae | XP_054047971.1 | 1132 |
| 11 | 131575974 | Poecile atricapillus (Black-capped chickadee) | Neognathae | XP_058688116.1 | 1125 |
| 11 | 102088152 | Columba livia (rock pigeon) | Neognathae | XP_005510283.1 | 1126 |
| 11 | 107310278 | Coturnix japonica (Japanese quail) | Neognathae | XP_032298647.1 | 1126 |
| 11 | 104461062 | Pterocles gutturalis | Neognathae | XP_010070812.1 | 1086 |
| 11 | 130146172 | Falco biarmicus (lanner falcon) | Neognathae | XP_056188019.1 | 1126 |
| 11 | 129202946 | Grus americana (Whooping crane) | Neognathae | XP_054672678.1 | 1028 |
| 12 | 119945562 | Tachyglossus aculeatus (Australian echidna) | Monotrema | XP_038622683.1 | 1116 |
| 12 | 100087634 | Ornithorhynchus anatinus (platypus) | Monotrema | XP_028924250.1 | 1079 |
| 13 | 127543775 | Antechinus flavipes (yellow-footed antechinus) | Metatheria | XP_051826012.1 | 1119 |
| 13 | 123231668 | Gracilinanus agilis (Agile Mouse Opossum) | Metatheria | XP_044513889.1 | 1123 |
| 13 | 122748454 | Dromiciops gliroides (monito del monte) | Metatheria | XP_043850025.1 | 1119 |
| 13 | 118837219 | Trichosurus vulpecula (common brushtail) | Metatheria | XP_036600048.1 | 1118 |
| 13 | 110197275 | Phascolarctos cinereus (koala) | Metatheria | XP_020826704.1 | 1119 |
| 13 | 100918272 | Sarcophilus harrisii (Tasmanian devil) | Metatheria | XP_031802128.1 | 1119 |
| 13 | 100028386 | Monodel. domestica (gray short-tailed opossum) | Metatheria | XP_007487041.2 | 1123 |
| 14 | 101356028 | Trichechus manatus latirostris (Florida manatee) | Sirenia | XP_012409888.2 | 1212 |
| 15 | 100654300 | Loxodonta africana (African savanna elephant) | Proboscidea | XP_003408268.1 | 1119 |
| 15 | 126058709 | Elephas maximus indicus (Indian elephant) | Proboscidea | XP_049709885.1 | 1119 |
| 16 | 103202460 | Orycteropus afer afer (Aardvarks) | Afroinsectiphilia | XP_007945544.1 | 1117 |
| 16 | 102862466 | Elephantulus edwardii (Cape elephant shrew) | Afroinsectiphilia | XP_006880602.1 | 1121 |
| 16 | 101648207 | Echinops telfairi (Small Madagascar hedgehog) | Afroinsectiphilia | XP_045146169.1 | 1091 |
| 17 | 137751018 | Eschrichtius robustus (Grey whale) | Cetacea | XP_068381580.1 | 1303 |
| 17 | 132351746 | Balaenoptera ricei (Rice's whale) | Cetacea | XP_059757710.1 | 1239 |
| 17 | 118883363 | Balaenoptera musculus (Blue whale) | Cetacea | XP_036685979.1 | 1295 |
| 17 | 103012702 | Balaenoptera acutorostrata (Minke whale) | Cetacea | XP_007185186.2 | 1314 |
| 18 | 130853881 | Hippopotamus amphibius kiboko | Ancodonta | XP_057592299.1 | 1117 |
| 19 | 505317 | Bos taurus (Cattle) | Ruminantia | XP_015329969.1 | 1119 |
| 19 | 101115717 | Ovis aries (Sheep) | Ruminantia | XP_027828841.2 | 1154 |
| 19 | 138088510, | Capricornis sumatraensis (Sumatran serow) | Ruminantia | XP_068839865.1 | 1119 |
| 19 | 122679320, | Cervus elaphus (Red deer) | Ruminantia | XP_043735850.1 | 1155 |
| 19 | 122451076 | Cervus canadensis | Ruminantia | XP_043339538.1 | 1155 |
| 19 | 129559976 | Moschus berezovskii (Chinese forest deer) | Ruminantia | XP_055287930.1 | 1119 |
| 20 | 100152934 | Sus scrofa (pig) | Suina | XP_001926150.2 | 1156 |
| 20 | 125129254 | Phacochoerus africanus (Common warthog) | Suina | XP_047639718.1 | 1140 |
| 21 | 123860208 | Mirounga angustirostris (Northern elephant seal) | pinnipedae | XP_045754950.1 | 1120 |
| 21 | 11801538 | Mirounga leonina (Southern elephant seal) | pinnipedae | XP_034868149.1 | 1120 |

| | | | | | |
|---|---|---|---|---|---|
| 21 | 118547638 | Halichoerus grypus (gray seal) | pinnipedae | XP_035967097.1 | 1119 |
| 21 | 116643255 | Phoca vitulina (harbor seal) | pinnipedae | XP_032277916.1 | 1120 |
| 21 | 114213789 | Eumetopias jubatus (Steller sea lion) | pinnipedae | XP_027964880.1 | 1120 |
| 21 | 113938568 | Zalophus californianus (California sea lion) | pinnipedae | XP_027479786.1 | 1120 |
| 21 | 112822942 | Callorhinus ursinus (northern fur seal) | pinnipedae | XP_025726311.1 | 1120 |
| 21 | 110578825 | Neomon. schauinslandi (Hawaiian monk seal) | pinnipedae | XP_021544050.1 | 1120 |
| 21 | 102730954 | Leptonychotes weddellii (Weddell seal) | pinnipedae | XP_006735615.1 | 1120 |
| 21 | 101374525 | Odobenus rosmarus divergens (Pacific walrus) | pinnipedae | XP_004408769.1 | 1120 |
| 22 | 132013148 | Mustela nigripes (Black-footed ferret) | Musteloidea | XP_059248862.1 | 1121 |
| 22 | 131826718 | Mustela lutreola (European mink) | Musteloidea | XP_059022217.1 | 1120 |
| 22 | 125099091 | Lutra lutra (Eurasian river otter) | Musteloidea | XP_047584393.1 | 1120 |
| 22 | 123955161 | Meles meles (Eurasian badger) | Musteloidea | XP_045882814.1 | 1120 |
| 22 | 122903939 | Neogale vison (American mink) | Musteloidea | XP_044100400.1 | 1120 |
| 22 | 116874564 | Lontra canadensis (N. Am. river otter) | Musteloidea | XP_032725145.1 | 1120 |
| 22 | 116575394 | Mustela erminea (ermine) | Musteloidea | XP_032172629.1 | 1120 |
| 22 | 101693793 | Mustela putorius furo (domestic ferret) | Musteloidea | XP_012916863.1 | 1120 |
| 23 | 100482362 | Ailuropoda melanoleuca (Giant panda) | Ursidae | XP_002922845.2 | 1120 |
| 23 | 123800478 | Ursus americanus (American black bear) | Ursidae | XP_045666152.1 | 1120 |
| 23 | 113253058 | Ursus arctos (Brown bear) | Ursidae | XP_026351684.1 | 1120 |
| 23 | 103681282 | Ursus maritimus (Polar bear) | Ursidae | XP_040481520.1 | 1120 |
| 24 | 101533744 | Ochotona princeps (American pika) | Lagomorpha | XP_004588168.2 | 1117 |
| 24 | 121166518 | Ochotona curzoniae (black-lipped pika) | Lagomorpha | XP_040851616.1 | 1117 |
| 24 | 133758382 | Lepus europaeus (European hare) | Lagomorpha | XP_062045531.1 | 1116 |
| 24 | 100341337 | Oryctolagus cuniculus (rabbit) | Lagomorpha | XP_008253877.2 | 1124 |
| 25 | 101967889 | Ictidomys tridecemlineatus (13-lined squirrel) | Rodentia (S) | XP_040149889.1 | 1121 |
| 25 | 107142282 | Marmota marmota marmota (Alpine marmot) | Rodentia (S) | XP_015341198.1 | 1121 |
| 25 | 113188789 | Urocitellus parryii (Arctic ground squirrel) | Rodentia (S) | XP_026253050.1 | 1121 |
| 25 | 114101557 | Marmota flaviventris (yellow-bellied marmot) | Rodentia (S) | XP_027802432.2 | 1121 |
| 25 | 124103886 | Marmota monax (woodchuck) | Rodentia (S) | XP_046316517.1 | 1121 |
| 25 | 124994774 | Sciurus carolinensis (gray squirrel) | Rodentia (S) | XP_047423001.1 | 1120 |
| 25 | 105994775 | Dipodomys ordii (Ord's kangaroo rat) | Rodentia (C) | XP_012883871.1 | 1117 |
| 25 | 109680115 | Castor canadensis (American beaver) | Rodentia (C) | XP_020010675.1 | 1117 |
| 25 | 122096329 | D. spectabilis (banner-tailed kangaroo rat) | Rodentia (C) | XP_042522821.1 | 1117 |
| 25 | 125360622 | P. longimembris pacificus (Pc. pocket mouse) | Rodentia (C) | XP_048214618.1 | 1116 |
| 25 | 101723077 | Heterocephalus glaber (naked mole-rat) | Rodentia (H) | XP_004842237.2 | 1120 |
| 25 | 102027018 | Chinchilla lanigera (long-tailed chinchilla) | Rodentia (H) | XP_005392155.1 | 1118 |
| 25 | 104851716 | Fukomys damarensis (Damara mole-rat) | Rodentia (H) | XP_033616456.1 | 1118 |
| 25 | 100526649 | Cavia porcellus (domestic guinea pig) | Rodentia (H) | NP_001185699.1 | 1111 |
| 25 | 110540668 | Meriones unguiculatus (Mongolian gerbil) | Rodentia (M) | XP_060240591.1 | 1120 |
| 25 | 126501455 | Alexandromys fortis (reed vole) | Rodentia (M) | XP_049998080.1 | 1116 |
| 25 | 127233320 | Phodopus roborovskii (desert hamster) | Rodentia (M) | XP_051055706.1 | 1124 |
| 25 | 127680472 | Apodemus sylvaticus (European woodmouse) | Rodentia (M) | XP_052031891.1 | 1125 |
| 25 | 129673566 | Psammomys obesus (fat sand rat) | Rodentia (M) | XP_055459399.1 | 1120 |
| 25 | 130887984 | Chionomys nivalis (European snow vole) | Rodentia (M) | XP_057646741.1 | 1116 |

| | | | | | |
|---|---|---|---|---|---|
| 25 | 31904594 | Peromyscus eremicus (cactus mouse) | Rodentia (M) | XP_059111663.1 | 1124 |
| 25 | 101834072 | Mesocricetus auratus (golden hamster) | Rodentia (M) | XP_021080916.1 | 1116 |
| 25 | 312896 | Rattus norvegicus (Norway rat) | Rodentia (M) | NP_997491.2 | 1125 |
| 25 | 277328 | Mus musculus (house mouse) | Rodentia (M) | NP_808449.1 | 1125 |
| 26 | 103585496 | Galeopterus variegatus (Sunda flying lemur) | dermoptera | XP_008564696.1 | 1119 |
| 26 | 134364008 | Cynocephalus volans (Philippine flying lemur) | dermoptera | XP_062935968.1 | 1119 |
| 26 | 105825356 | Propithecus coquereli (Coquerel's sifaka) | primate | XP_012518560.1 | 1090 |
| 26 | 100942247 | Otolemur garnettii (small-eared galago) | primate | XP_012666997.1 | 1110 |
| 26 | 123644510 | Lemur catta (Ring-tailed lemur) | primate | XP_045416598.1 | 1136 |
| 26 | 117078803 | Trachypithecus francoisi (Francois's langur) | primate | XP_033058185.1 | 1116 |
| 26 | 116813194 | Hylobates moloch (silvery gibbon) | primate | XP_032615472.1 | 1118 |
| 26 | 126961477 | Macaca thibetana thibetana | primate | XP_050657875.1 | 1119 |
| 26 | 128563615 | Nycticebus coucang (slow loris) | primate | XP_053415000.1 | 1146 |
| 26 | 464230 | Pan troglodytes (chimpanzee) | primate | XP_519806.3 | 1119 |
| 26 | 129042811 | Pongo pygmaeus (Bornean orangutan) | primate | XP_054355101.1 | 1119 |
| 26 | 129493029 | Symphalangus syndactylus (siamang) | primate | XP_055154638.1 | 1119 |
| 26 | 694623 | Macaca mulatta (Rhesus monkey) | primate | XP_001083172.2 | 1119 |
| 26 | 8989 | Homo sapiens (human) | primate | NP_015628.2 | 1119 |
